# Dyrk1b is a key Regulatory Kinase Integrating Fgf, Shh and mTORC1 signaling in Skeletal Muscle Development and Homeostasis

**DOI:** 10.1101/2020.04.22.055947

**Authors:** Neha Bhat, Anand Narayanan, Mohsen Fathzadeh, Anup Srivastava, Arya Mani

**Affiliations:** Cardiovascular Research Center, Department of Internal Medicine, Yale University School of Medicine, New Haven, CT, 06520, USA; Department of Genetics, Yale School of Medicine, New Haven, CT, 06511; Psychiatry and Behavioral Sciences, Stanford University, Palo Alto, CA, 94305

**Author notes:** Authors have contributed equally to the manuscript (Neha Bhat, Anand Narayanan). Correspondence to: Arya Mani, Yale Cardiovascular Research Center, 300 George Street, New Haven, USA 06511.

**Keywords:** *dyrk1b*, *myoD*, myogenesis, sarcopenic obesity, Shh, Fgf, human genetics, fate determination, inherited metabolic syndrome

## Abstract

The advent of human genetics has provided unprecedented opportunities for discovery of novel disease pathways. Mutations in *DYRK1B* have been associated with metabolic syndrome and sarcopenic obesity in humans, underscoring the critical role of the encoded protein in skeletal muscle development and homeostasis. By the novel creation of Dyrk1b knockout zebrafish models we demonstrate that Dyrk1b kinase activity is critical for specification of the paraxial *myoD*. Mechanistically, Dyrk1b mediates and amplifies Fgf signaling in the paraxial domain by the transcriptional suppression of its negative feedback inhibitor *sprouty1*. In the adaxial *myoD* domain, Dyrk1b amplifies Shh signaling and partially rescues defects caused by its disruption. The investigations of C2C12 terminal differentiation revealed that Dyrk1b also plays a critical role in myofiber fusion. Combined biochemical and proteomic analysis of C2C12 myoblasts undergoing differentiation showed that Dyrk1b kinase activation is induced by shh inhibition, and triggers differentiation by inhibiting mTOR, subsequent upregulation of 4e-bp1 and induction of autophagy. In conclusion, we demonstrate that Dyrk1b plays a critical role in sustaining myocyte specification and differentiation by integrating Fgf, Shh and mTORC1 signaling pathways.

## INTRODUCTION

The syndrome of Sarcopenic Obesity (SO), characterized by a combination of truncal obesity and reduced skeletal muscle mass is an emerging risk factor for the metabolic syndrome (MetS), (1) and is a greater risk for cardiovascular diseases, and type2 diabetes than obesity alone. Accordingly, the burden of type 2 diabetes is most rapidly increasing in populations with low to normal body mass index but high prevalence of sarcopenic obesity (2, 3). The rapid trend of aging population and the growing sedentary life styles are other impetus for the recent surge in sarcopenia (4), which has prompted universal efforts to reduce its burden. Unfortunately, strategies to improve skeletal muscle mass have been largely unsuccessful due to the limited understanding of myogenesis as an integrated process in humans.

SO has strong genetic components that are linked to the risk for type2 diabetes and atherosclerosis (5-9). The advent of human genome project has provided an excellent opportunity for the discovery of novel disease genes and their cognate pathways. Specifically, major progress has been made in genetic studies of single gene disorders. This in turn has led to the discovery of key driver nodes of complex biological networks involved in development and disease. Most recently, independent, rare and novel genetic variants with large effect sizes in *DYRK1B* gene have been shown to underlie atherosclerosis, MetS and sarcopenic obesity by using whole exome sequencing (10). This discovery provided a unique opportunity to investigate developmental and disease pathways of the human skeletal muscle.

DYRK1B protein is a dual-specificity kinase, with the tyrosine phosphorylation primarily limited to auto-phosphorylation (Y271/273) and an arginine-directed serine/threonine kinase activity towards its substrates (10, 11). It is a pancellular kinase (9, 12), that is highly expressed in the skeletal muscle and has been reported to promote *in vitro* differentiation of myofibers in C2C12 myoblast cells (13-15). The mechanisms by which Dyrk1b promote myofiber differentiation are relatively unexplored. Surprisingly, no *in vivo* genetic model of DYRK1B and its characterization has been reported to this date. Consequently, its roles during skeletal muscle specification, prior to myoblast differentiation have never been explored. We embarked on investigating its role during early myogenesis in zebrafish due to the ease of genetic engineering, external fertilization and the transparency of embryonic development. Although, the skeletal muscle in zebrafish is predominantly made of fast twitch fibers (16, 17), the fundamental pathways involved in regulation of myogenic specification are highly conserved across metazoans (18).

The myogenesis across metazoans is regulated by the transcription factors (TF) called Muscle regulatory factors (MRFs) which includes MyoD, Myf5, Myogenin and MRF proteins (19, 20). The basic helix-loop-helix TF MyoD is the central protein sufficient to orchestrate differentiation of numerous cell types to skeletal muscle fate (21, 22). The MRFs cooperate with the coregulatory proteins of Mef2 class of transcription factors to stimulate myofiber differentiation (23). During zebrafish embryonic myogenesis, *myf5, myoD* and *myogenin* are co-expressed in cells adjacent to the axial mesoderm (aka. adaxial *myoD*) which differentiate into the slow muscle fibers (Wolff et al., 2003, Reifers et al., 1998). In the paraxial myotomal somitic domain, *myoD* (aka. paraxial *myoD*) and *myogenin* are upregulated as somitogenesis proceeds (17) and these cells differentiate into fast-twitch fibers. Based on earlier *in vivo* studies, the adaxial and paraxial *myoD* expression is regulated by distinct signaling pathways: Shh and Fgf, respectively (24). In zebrafish, the knockout of Shh canonical receptor *smoothened* results in loss of adaxial *myoD* with no detectable effect on paraxial *myoD* (25). Conversely, loss of Fgf8 leads to complete loss of the paraxial *myoD* with no change reported in the adaxial domain (Groves et al., 2005). Thus, Shh and Fgf act in a mutually exclusive but coordinated fashion to drive expression of *myoD* in these distinct domains within the somites. However, the downstream effectors of Shh and Fgf pathways during muscle specification are largely unknown.

During the subsequent stage of differentiation of myoblasts into myofibers, Fgf (26) and Shh (27) signaling pathways play much more different, complex and dynamic roles. While data from some investigators supports the role of Fgf signaling in myotube fusion (28, 29)other studies have either contradicted these findings(30, 31) or have suggested stage specific function of FGF during myotube differentiation (32, 33) (34). Similarly, opposite results for the role of Shh on myoblast fusion has been reported (27, 35). These discrepancies could be due to the variability of the dosage, timing and duration of its pharmacological manipulations, necessitating more in-depth investigations. In addition, pathways downstream from DYRK1B in regulation of myogenesis were relatively unknown.

We comprehensively investigated the role of Dyrk1b kinase in myogenic specification, and differentiation and examined its role in integrating different myogenic pathways into a coordinated developmental process. We generated a CRISPR/Cas9-mediated Dyrk1b zebrafish knockout model. We found that this kinase is indispensable for survival as Dyrk1b-/- embryos did not survive beyond 72hpf. We subsequently examined the role of Dyrk1b in muscle specification and differentiation in the context of pharmacological and genetic perturbation of Shh and Fgf pathways and delineated its role in 1) regulation of paraxial and adaxial domains of *myoD* expression in zebrafish, and in 2) myofiber differentiation in mouse C2C12 cells. We find that in zebrafish myogenic progenitors and C2C12 myoblasts, Dyrk1b interact with Fgf and Shh signaling in strikingly contrasting ways. In zebrafish, Dyrk1b acts downstream of the Fgf signaling to specify paraxial *myoD* domain and increases Shh signaling to specify adaxial *myoD* cells. In C2C12 cells, Dyrk1b kinase activation is enhanced by Shh inhibition, which results in myofiber differentiation. Using proteomics analysis in C2C12 cells, we identified genes regulated by DYRK1B during myofiber differentiation and uncovered a network of co-regulated pathways that are activated upon Dyrk1b inhibition of mTORC1 and are associated with autophagy. Strikingly, the induction of excessive autophagy by DYRK1B^R102C^ mutation was coupled with impaired myogenesis in C2C12 cells. Altogether, we demonstrate the role of Dyrk1b as a key node integrating Fgf, Shh, and mTORC1 signaling during myogenesis.

## RESULTS

### Dyrk1b is expressed in the somitic mesoderm, that harbors the precursors of skeletal muscles

We chose zebrafish as a vertebrate model to study the dynamic process of skeletal muscle development due to the ease in creating genomic alterations, the external fertilization, the transparency of the embryos and the high levels of sequence identity (80%) and similarity (94%) in the kinase domains of zebrafish and human Dyrk1b. We first examined the expression of *dyrk1b* in the zebrafish by whole mount *in-situ* hybridization (WISH), using two digoxigenin-labeled anti-sense probes and a control sense RNA probe. We performed WISH at various stages of embryogenesis starting from single cell stage to 24 hpf. Examination of mRNA expression of *dyrk1b* revealed that transcripts are deposited maternally, and the gene is expressed from 1-cell stage through 4 and 8-cell stages (Fig. 1A-D, n=30 embryos at each stage). After the onset of zygotic transcription, the expression is broadly seen throughout the embryo at 9hpf (Fig. 1E), both ventrally and dorsally. At mid-somitogenesis (14 hpf), the transcripts are expressed in the entire dorsal region, but were excluded from the posterior axial domain of the embryo (red asterisk, Fig. 1F, n=30 embryos at each stage). At 24-hpf stage, the *dyrk1b* transcripts are localized in the anterior neuroectoderm including the forebrain, midbrain and hindbrain (Fig.1G, n=30 embryos at each stage). The cross sections in the anterior embryonic domains revealed expression in the forebrain and eyes (Fig. 1H) at 24 hpf. In the posterior domain, *dyrk1b* is expressed in mesodermally derived somites (red asterisks, in Fig. 1I), but excluded from the notochord (red asterisk in Fig. 1G and red arrow in Fig. 1I) and posterior neural tube (Fig.1I NT, sections were examined at 100um intervals for n=5 embryos for each stage). The sense-control probe showed no staining (Fig. 1K) when compared to the antisense probe (Fig. 1J). The second anti-sense probe generated against *dyrk1b*, showed an identical expression pattern by WISH (SFig.1).

**Figure 1.**
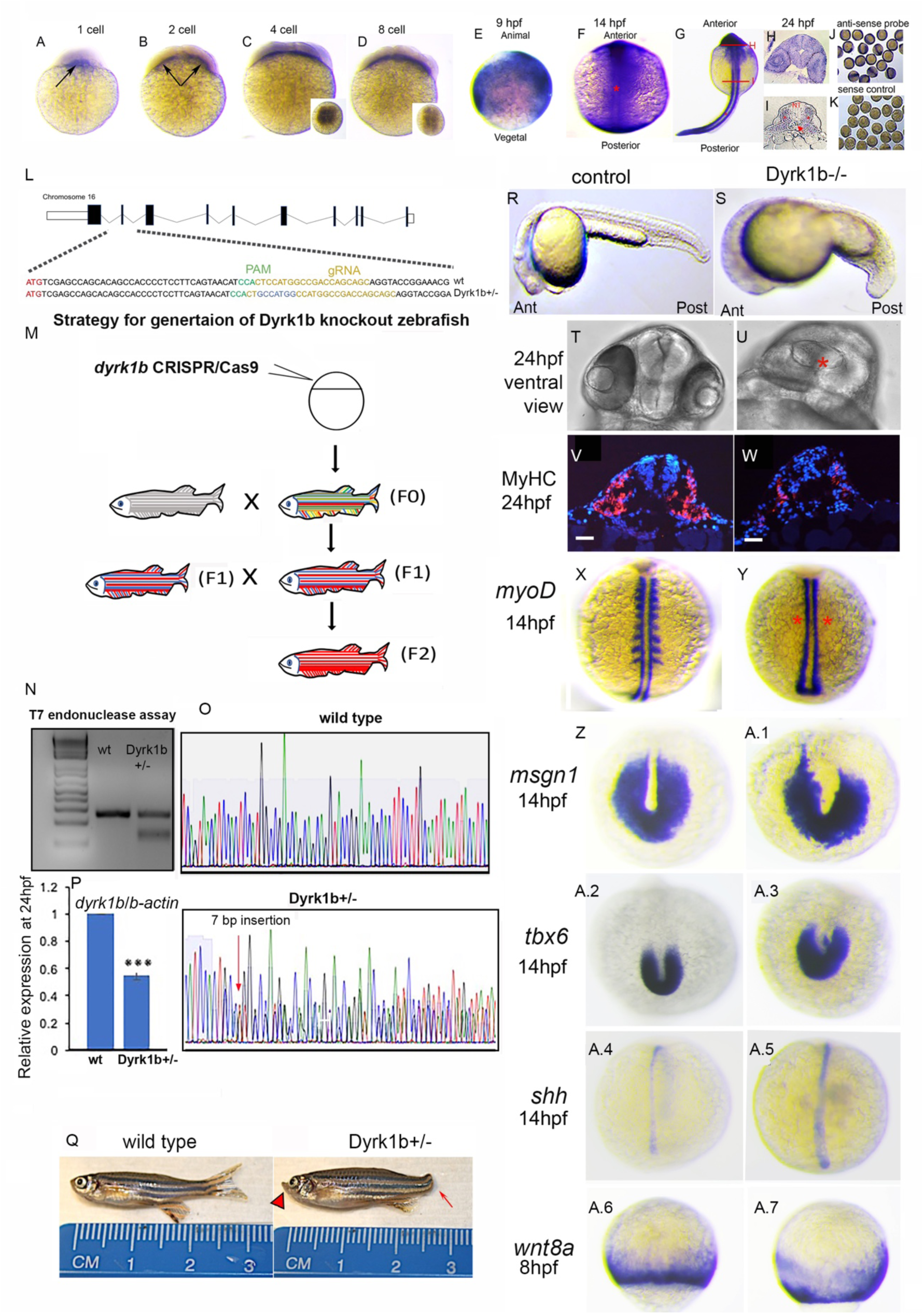
Knockdown of *dyrk1b* in the zebrafish causes loss of paraxial myoD, the central regulator of embryonic myogenesis. **A-I** WISH for *dyrk1b* transcripts using an anti-sense probe at different stages of zebrafish embryonic development as indicated. Arrows on A and B indicate transcripts at one and two cell stages, respectively. Insets on C and D show animal pole view. A-E lateral views, F-G dorsal views, H and I are cross-sections at regions indicated by red lines on G. Red asterisks on F and G indicate the unstained axial domain. Red asterisks on I denote somites, red arrow denotes notochord, NT=neural tube. **J**,**K** are group shots of zebrafish embryos stained with anti-sense and the control sense probe. **L.** Schematic diagram of the *dyrk1b* locus in the zebrafish genome. Boxes indicate the exons (not drawn to scale). The dotted lines from second exon indicate the translation start site (in red). Guide RNA is highlighted in yellow and PAM region in green. wildtype and *dyrk1b+/-* sequence are shown, 7bp insertion induced by CRISPR/Cas9 is highlighted in blue. **M** Schematic showing Dyrk1b CRISPR/Cas9 injection and transmission of induced mutation. **N** T7 assay with one band for Wildtype and two bands for *dyrk1b+/-*, indicating the heteroduplex formed by CRISPR/Cas9 activity. **O.** Sanger sequencing images of amplified segments of wildtype and *dyrk1b+/-*, demonstrating the site of 7bp insertion (red arrow) and the ensuing frame shift. **P.** *dyrk1b* mRNA expression levels (mean ± s.e.m) in the wildtype and Dyrk1b+/- zebrafish embryo at 24hpf (*** p≤ 0.001, Students ttest, 2 tailed, n=30 pooled embryos). **Q** Micrographs of 3-month-old adult heterozygous Dyrk1b+/- fish showing lack of caudal fin (red arrow) and jaw malformation (red arrowhead). **R, S** Lateral views of wildtype and Dyrk1b-/- embryos at 24hpf showing shortened body axis and defective somites in Dyrk1b-/- fish. **T, U** Ventral views of 24 hpf wildtype and Dyrk1b-/- embryos, red asterisk in R indicates cyclopic eyes. **V, W** Immuno-fluorescent staining (IF) for Myosin Heavy Chain1 on cross-section of 24hpf Dyrk1b-/- and control embryos. **X, Y** WISH for *myoD* at 14hpf in *dyrk1b* CRISPR/Cas9-injected fish, red asterisks in Y indicate loss of paraxial *myoD* (dorsal views). **Z-A.7** WISH for different markers of genes in the myogenic pathway, as indicated. Z-A.3 tail bud view at 14hpf, A.4-A.5 dorsal views, 14 hpf, A.6-A.7 lateral views at ∼80% epiboly, *indicates p≤0.05, ** p≤ 0.01, Students ttest, 2 tailed. Scale bars indicate 200 µm.

**Figure S1.**
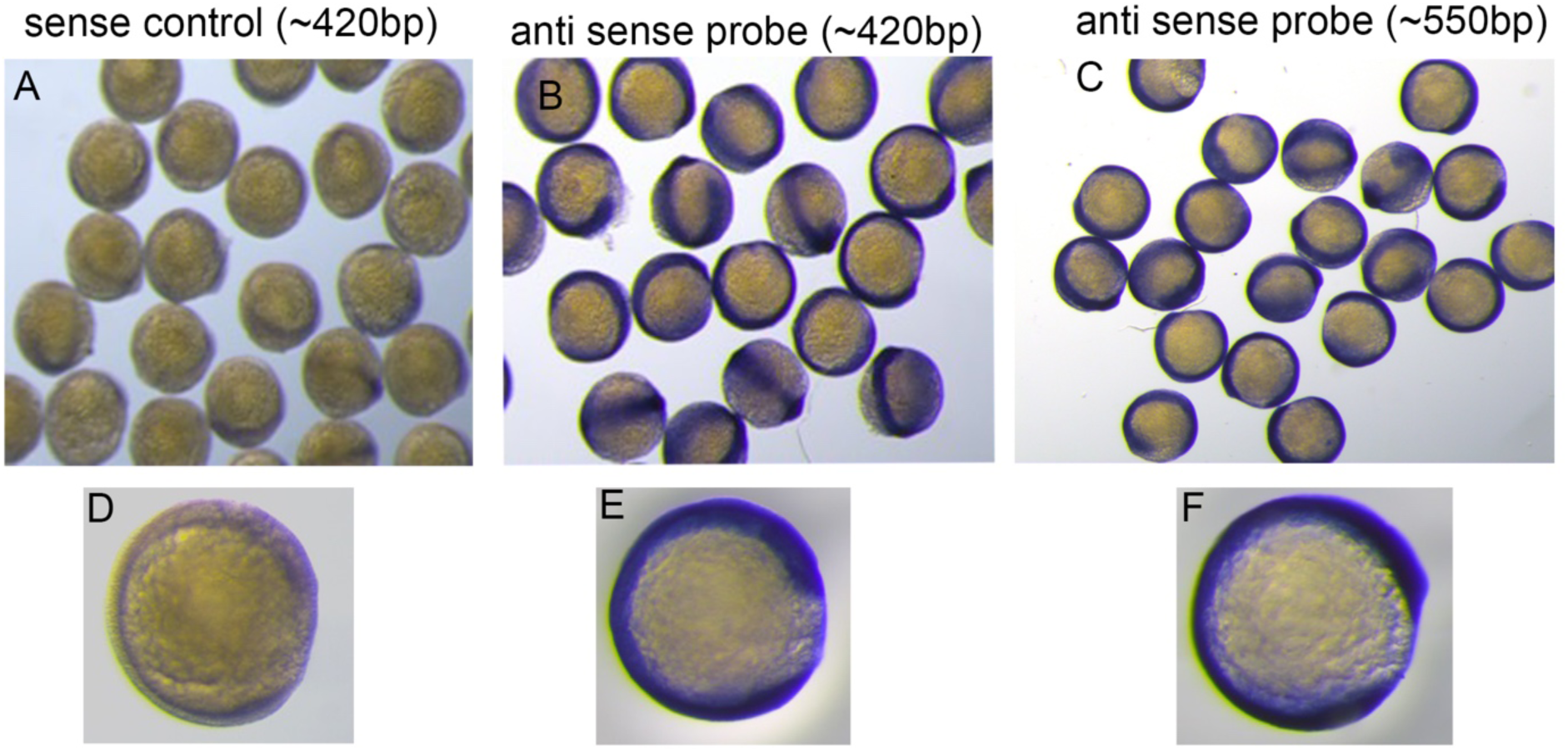
WISH of two anti-sense probes for Dyrk1b exhibiting identical expression pattern A-C Group shots of 14hpf embryos revealing Dyrk1b transcripts by WISH. D-F individual embryos from the respective in-situ (lateral view).

### Dyrk1b is required for normal myogenesis, brain and eye development in zebrafish

To determine the function of *dyrk1b* during zebrafish development, we employed CRISPR/Cas9 mediated gene editing to create stable knockouts (36). The major caveats of *dyrk1b* knockdown, previously attempted using anti-sense morpholino (37) includes widely reported off-target effects of the morpholinos and the lack of comparison of the observed phenotypes with a stable knockout (38). In this study, we generated stable *dyrk1b* knockout zebrafish, using two different guide RNAs (gRNA): one targeting the translation start site in exon 2 (Fig.1L, SFig.2A, for gRNA design and off-target analysis see the Materials and Methods) and another gRNA targeting the kinase domain of the *dyrk1b* gene (SFig.2B). Both gRNAs successfully and independently targeted the *dyrk1b* gene, generated indels and resulted in identical phenotypic traits. We employed three different methods to confirm the efficient disruption of the *dyrk1b* gene in the random clutch of wild type zebrafish embryos. First, 6 FAM (Fluorescein amidite) labeled primers were used, which revealed insertions and deletions (indels) created by the gRNA activity. Second, the T7 endonuclease activity assay was performed to further confirm the presence of indels and final confirmation was obtained by sanger sequencing. These different steps will be discussed again below for generation of stable F2 zebrafish knockouts. The F0 embryos showed the following consistent and striking phenotypes at 24hpf: cyclopic eyes, shortened and curved body axis. Nearly 30% of embryos were lethal. We raised the remaining ca. seventy percent surviving embryos to adulthood and screened for the founder fish (F0). The F0 fish, were inherently mosaic carrying several different *dyrk1b* mutations in different cells (indicated by a multicolored fish in schematic Fig. 1M) and were crossed to wild-type to generate F1 fish (Fig. 1M). After confirming the successful transmission of the mutation by T7 assay (Fig. 1N), the F1 fish were sequenced to identify the nature of the mutation. We identified three independent mutations in the *dyrk1b* gene in F1 fish (SFig. 2A). For the purpose of the study, we focused on the carrier fish that had an insertion of 7 bp near the translation start site of *dyrk1b* gene (Fig 1O). The total length of Dyrk1b is 751 amino acids, and this insertion caused a premature stop codon at the codon 59. The *dyrk1b* mRNA levels in the heterozygotes were about 50% of the wild type siblings as expected (Fig. 1P). The F1 heterozygote fish carrying this 7 bp insertion were intercrossed to obtain the F2 generation (schematic Fig. 1M). The adult heterozygote F1 fish showed protuberance of lower jaw, complete loss of the caudal fin and exhibited swimming defects (Fig.1Q).The F2 homozygous embryos showed similar traits as observed in F0 generation at 24hpf: shortened and curved body axis (Fig.1 R, S), visible loss of somites, cyclopic eyes (Fig. 1T, U) and were embryonic lethal and did not survive beyond 72 hpf.

**Figure S2.**
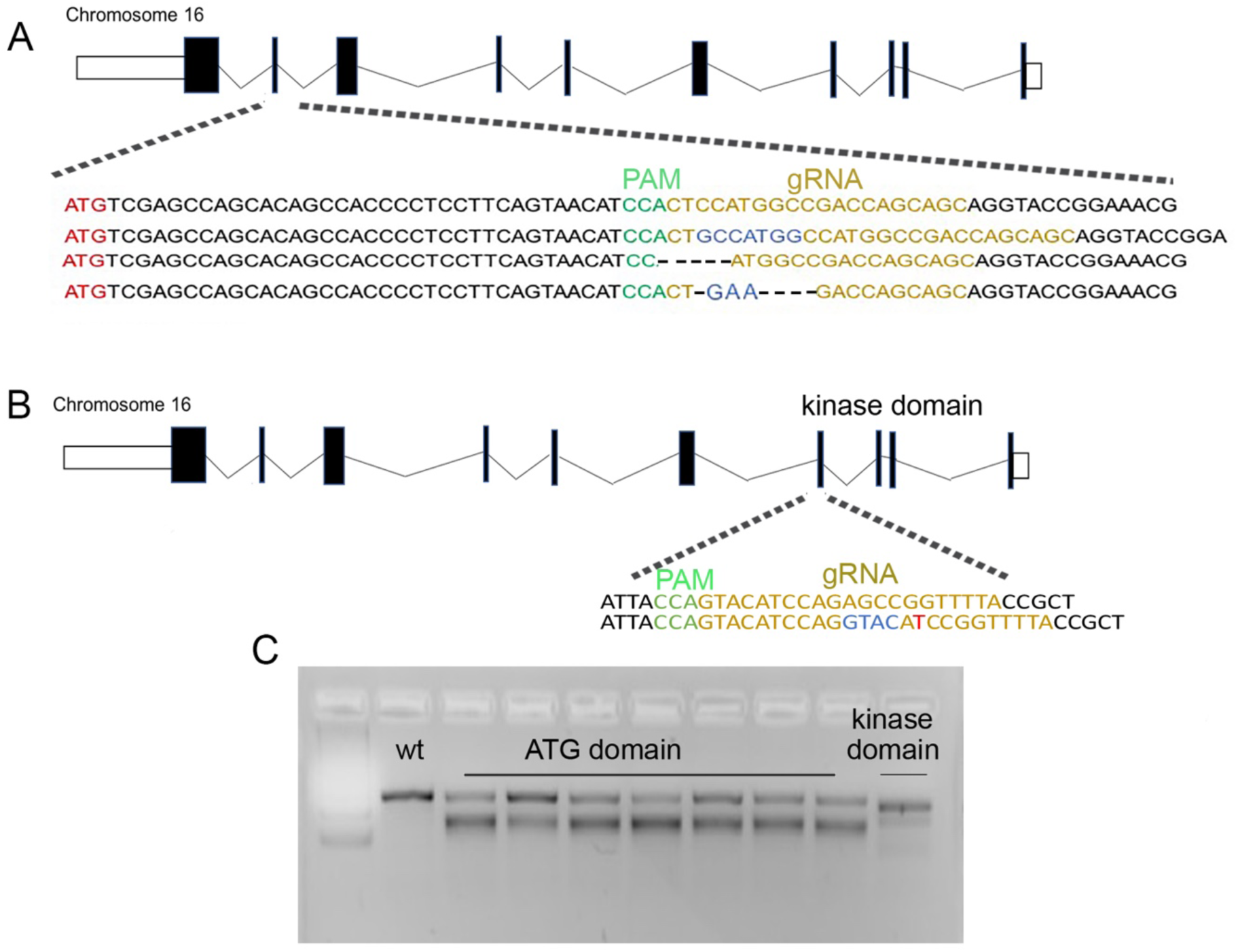
Reference DNA sequence flanking translation start site and kinase domain and the sequences of 3 mutations generated using CRISPR/Cas9. **A.** A schematic diagram of the *dyrk1b* locus in the zebrafish genome. Black boxes indicate exons (not drawn to scale). Dotted lines from the second exon indicate the translation start site (in red). Guide RNA is highlighted in yellow and PAM region is in green. The reference sequence is shown on the top and the 3 mutations are shown underneath **B.** CRISPR/Cas9 targeting the kinase domain creates a 4bp insertion and one-base-pair change (in red). **C.** T7 assay with one band for Wildtype (WT) and two bands for *dyrk1b+/-*, indicating the heteroduplex formed by CRISPR/Cas9 activity targeting exons carrying translation start site and kinase domain respectively.

### Dyrk1b regulates *myoD* expression in paraxial mesoderm

The expression of *dyrk1b* in the somites (Fig. 1I), and the shortened body axis (Fig. 1R, S) of Dyrk1b homozygous knockout F2 embryos at 24hpf, prompted the examination of skeletal muscle fibers, that are specified from the myotomal segments of somites (39). Immunofluorescent (IF) staining for Myosin heavy chain1 (MyHC1) in the Dyrk1b^-/-^ F2 embryos revealed a drastic reduction of myofibers at 24hpf (Fig.1 V, W, n=10 embryos of each genotype were sectioned, and cross sections in the trunk region were examined at intervals of 100um per embryo with identical results). To characterize if the loss of myofibers results from the molecular events occurring at earlier stages, we performed WISH for several unique but partially overlapping myogenic markers starting from mid-gastrulation through somitogenesis. This investigation reproducibly revealed complete loss of myogenic differentiation (*myoD)* expression in the paraxial mesoderm in F2 Dyrk1b -/- embryos (Fig.1X, Y, n=50 in each group for each experiment, n=3 biological replicates). Based on three independent breeding experiments, an average of 26.3 embryos from a clutch of total 99.3 F2 embryos were homozygous (Dyrk1b^-/-^) and showed complete loss of paraxial *myoD* at 14 hpf (bilateral myoD loss in 1Y and Table S1). Next, we examined expression of mesogenin1 (*msg1*), myogenic factor 5 (*myf5*), myogenin1 (*mygn1*) and T-box transcription factor 6 (*tbx6*). *mesogenin* (*msg1*), expressed in the posterior paraxial presomitic mesodermal progenitors (40), *tbx6*, expressed bilaterally in the somitic precursors anterior to *msg1* (41) and *myogenin*, expressed primarily in adaxial muscle precursors showed either no change or incremental expansion in the expression domains, in the Dyrk1b-/-vs. wildtype embryos at 14hpf (Fig.1Z-A.3, SFig.3A, n=50 embryos for each genotype, from two independent experiments). These expression patterns suggest that somitic progenitors are specified normally in the absence of Dyrk1b (39). Next, we examined the notochord marker, *shh* (42) and *wnt8a*, a marker of early mesodermal patterning (43) which showed no major changes in the Dyrk1b-/- embryos at the end of gastrulation (Fig 1A.4-A.7, n=50 embryos for each genotype, from two independent experiments). Interestingly, myogenic factor 5 (*myf5)* expressed in the myoblasts and the posteriorly localized somitic progenitors was upregulated in Dyrk1b-/- vs. wildtype embryos (SFig.3A, n=50 embryos for each genotype, from two independent experiments). As we will discuss later, the expansion in the *myf5* expression domain was also observed upon loss of Fgf signaling (see below). Altogether, these data reveal that Dyrk1b is dispensable for the specification of the presomitic progenitors but plays an essential function at the onset of *myoD* expression, the latter being the central regulator of embryonic and adult myogenesis.

### The kinase activity of Dyrk1b is essential for paraxial *myoD* specification

We next examined whether kinase activity of Dyrk1b is required for specification of *myoD* positive cells in zebrafish. To this end, we misexpressed human DYRK1B in zebrafish. We noted about 3-fold increase in *dyrk1b* transcripts upon its overexpression (Fig.2A). The injection of wildtype *DYRK1B* mRNA into wild-type zebrafish at one-cell stage extended the expression of *myoD* in each somite in the anterior-posterior axis. (Fig. 2B, C). To test whether kinase-activity of Dyrk1b was necessary, we mutated two residues in Dyrk1b: tyrosine 273 residue (Y273F) that undergoes cis-auto phosphorylation and the Lysine 140 in the ATP binding region of the catalytic domain (K140R) (10, 11). The *DYRK1B*^*Y273F, K140R*^ mRNA overexpression failed to expand paraxial *myoD* domain, indicating that kinase activity of Dyrk1b is important for myogenic specification (Fig. 2D, red arrow) and may have a dominant negative effect as suggested before (44).

**Figure S3.**
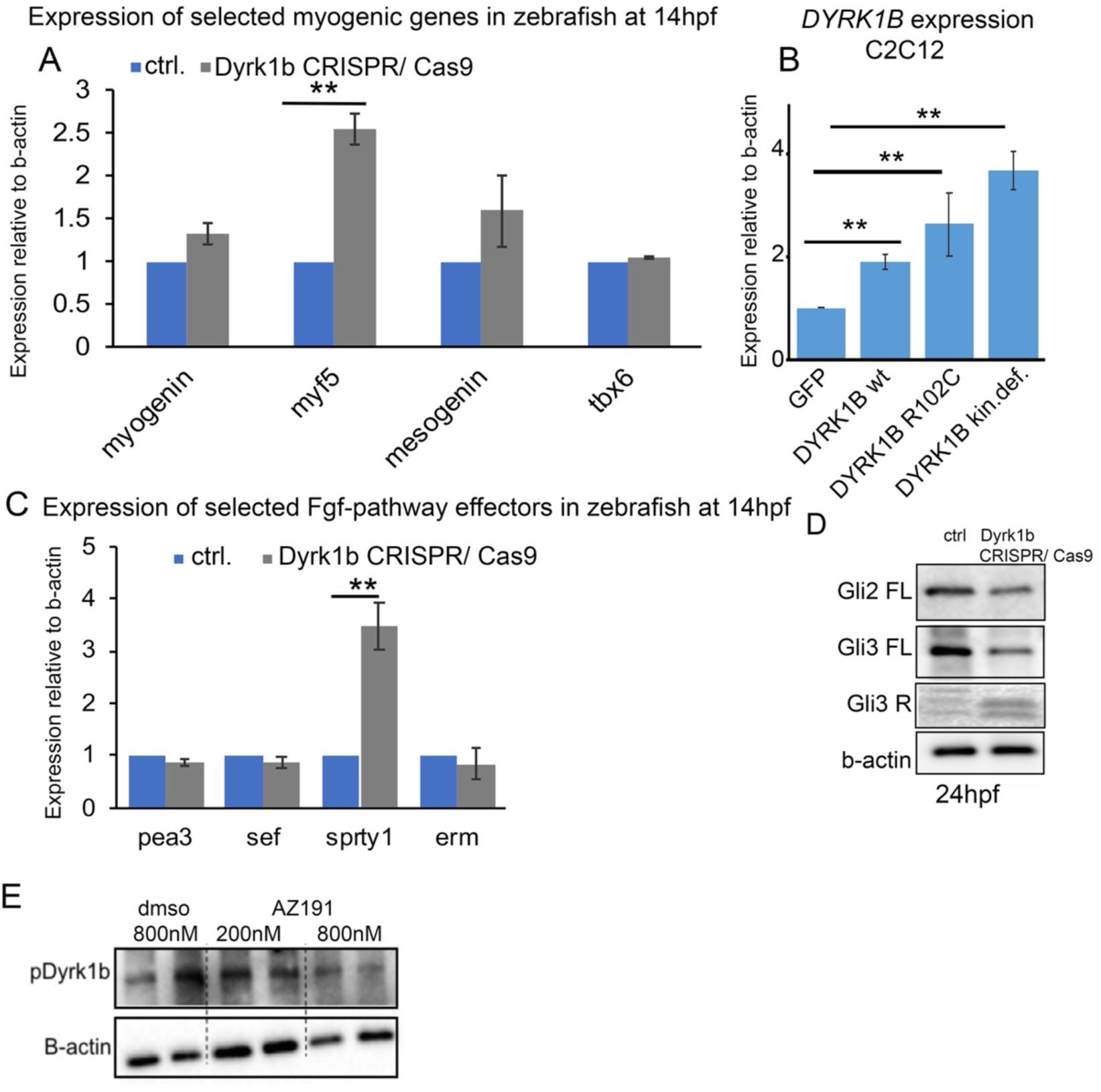
**A** Expression by RT-qPCR of selected myogenic differentiation factors in Dyrk1b knockout zebrafish embryos at 14hpf. **B** RT-qPCR showing Dyrk1b levels in the C2C12 cells transfected with the indicated constructs. **C** RT-qPCR of the indicated genes in the Fgf signaling pathway in Dyrk1b knockout zebrafish embryos. **D** The expression of Shh effectors Gli2 and Gli3 in Dyrk1b knockout embryos compared to control zebrafish embryos at 24hpf. **E** The expression of pDyrk1b, using pHIPK2 antibody (45) in primary cells treated with different concentrations of dmso or AZ191.

**Figure 2.**
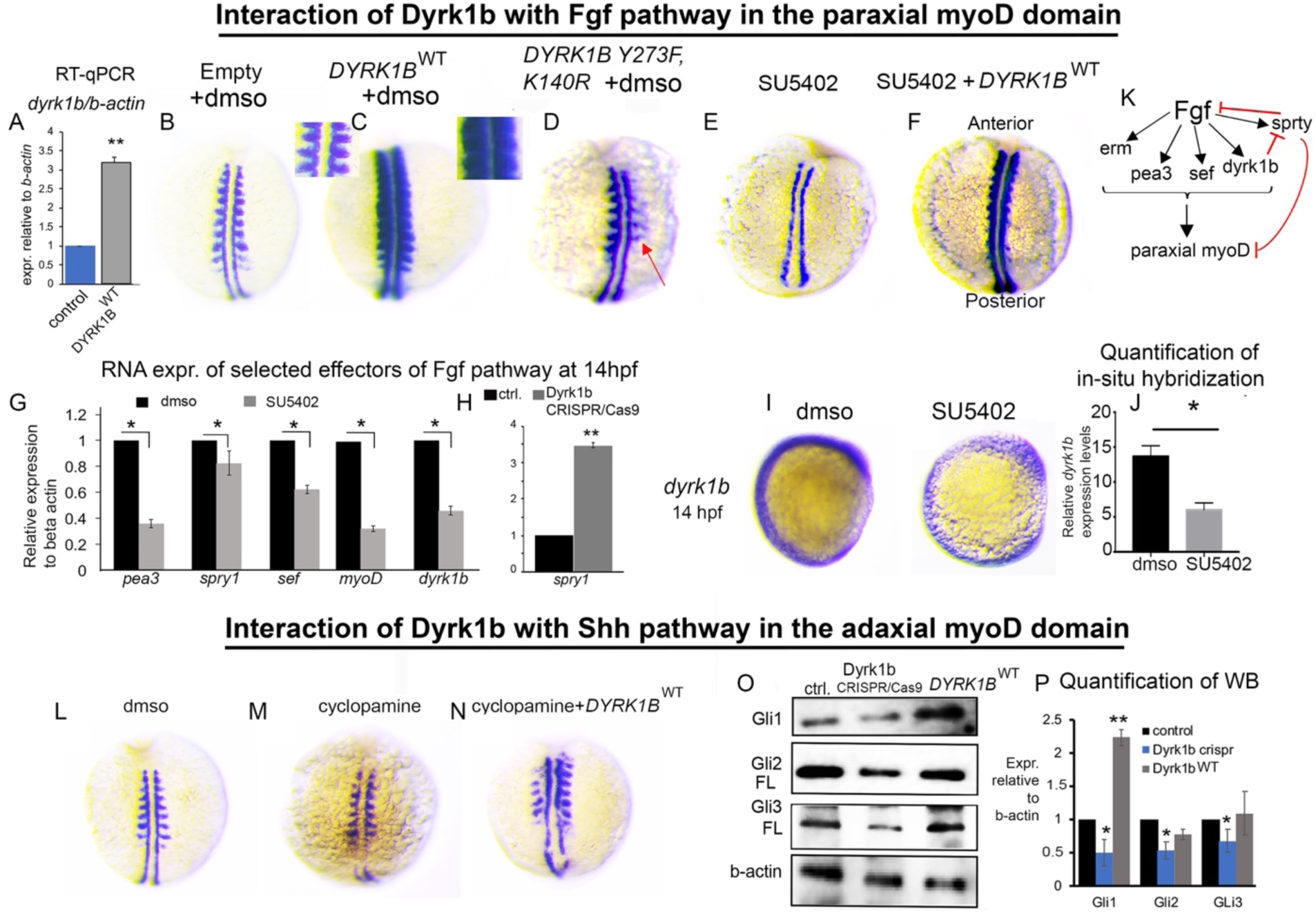
Dyrk1b mediates Fgf and Shh signaling in the paraxial and adaxial expression domains respectively. **A** *dyrk1b*, normalized to b-actin, is elevated in zebrafish overexpressing human wildtype *DYRK1B*. **B-D** WISH for *myoD* (dorsal views) in wild type embryos injected with plasmids containing no cDNA (empty), human wildtype *DYRK1B*, and kinase-defective (Y273F, K140R), and exposed to dmso, at the same concentration at SU5402, at 14hpf. Inset in B and C are magnifications of *myoD* expression for better visualization. **E, F** WISH for *myoD*, showing loss of paraxial *myoD* after treatment FGF inhibitor SU5402 (E) and its rescue by *DYRK1B* overexpression in SU5402 treated embryos at 14hpf. **G** mRNA expression of Fgf syn-expression genes, *myoD* and *dyrk1b* in dmso or SU5402 treated embryos at 14hpf. SU5402 was added at the end of gastrulation. **H** mRNA expression of *sprouty1* in *dyrk1b* knockout embryos at 14hpf. **I, J** WISH and quantification (by Image J) for *dyrk1b* in SU5402 treated embryos. **K** the model depicting that Dyrk1b mediates Fgf signaling and also amplifies it by suppressing *sprouty1*, the feed-back inhibitor of the Fgf pathway. **L-M** WISH(dorsal views) for *myoD* in 14hpf embryos, showing loss of adaxial myoD in cyclopamine treated embryos **N** Rescue of adaxial *myoD* by human *DYRK1B* overexpression in cyclopamine treated embryos. **O, P** Western blot images and their quantification (mean ± s.e.m) showing the expression of Shh effectors Gli1, Gli2 and Gli3 in control wild type embryos, Dyrk1b knockout and overexpressing zebrafish embryos at 14hpf. B-F, L-N dorsal views, I lateral views *indicates p ≤0.05, ** p≤ 0.01, *** p≤ 0.001, Students ttest, 2 tailed.

### Dyrk1b mediates Fgf signaling of the paraxial *myoD* domain in zebrafish

Paraxial *myoD* was previously reported to be transcriptionally regulated by fibroblast growth factor (Fgf) signaling (24). Fgf8 mutant embryos (*acerebellar*) as well as embryos treated with FgfR-specific tyrosine kinase inhibitor SU5402 show complete loss of *myoD* expression in the paraxial somites (24). Since Dyrk1b-/- embryos phenocopy the Fgf mutants’ loss of *myoD* in the paraxial domain (Fig. 1X, Y) we decided to explore the epistatic interaction between Fgf and Dyrk1b in zebrafish. Note, that *myf5* expression domains are expanded in Fgf8 mutants and SU5402 treated embryos (24) similar to the Dyrk1b-/- embryos (SFig.3A). We inhibited Fgf signaling by administrating SU5402 to the embryos post gastrulation to avoid potential interference from effects of Fgf disruption during gastrulation on subsequent myogenesis (24). The embryos treated with SU5402 showed loss of *myoD* in paraxial domain and recapitulated *acerebellar* mutants (Fig. 2E) when compared to dmso control (Fig. 2B). To determine whether Dyrk1b mediates Fgf signaling during paraxial *myoD* specification, DYRK1B^WT^ was overexpressed at one-cell stage and treated with SU5402 post-gastrulation. Strikingly, the entire domain of paraxial *myoD* was rescued in this condition (Fig. 2F) (86.3% ± 3.8% embryos were rescued; P-value= 0.001, average 35± 10 embryos per experiment, three independent experiments). This finding demonstrates the critical role of Dyrk1b in mediating Fgf signaling during specification of paraxial *myoD*. Next, we examined whether Fgf signaling regulates *dyrk1b* transcription as a mechanism for specification of paraxial *myoD*. The expression levels of Fgf8 synexpression group genes *pea3, sprty1*, and *sef* (46, 47) and the expression of *myoD* were reduced, ensuring a sufficient blockage of the Fgf signaling pathway with SU5402 (Fig. 2G). Strikingly, the inhibition of Fgf led to 50% reduction in the *dyrk1b* expression levels, as assessed by both RT-qPCR (Fig. 2G) and WISH (Fig. 2 I, J). These data indicate that *dyrk1b* mediates Fgf signaling in the paraxial *myoD* domain. Next, we investigated whether Dyrk1b mediates Fgf signaling in the paraxial domain by regulating expression of Fgf effector genes. The positive regulators of Fgf signaling *pea3, sef* and *erm* remained unchanged (SFig.3C) while the negative regulator of Fgf signaling, *spry1* was significantly upregulated in Dyrk1b-/- embryos (Fig.2H). This finding is consistent with a previous study that demonstrated the expansion of paraxial *myoD* domain by disruption of *spry4* in zebrafish (48), indicating a stringent negative feedback mechanism on Fgf signaling to spatially limit the paraxial *myoD* domain (48). Altogether, these data suggest the following model: Fgf signaling in the paraxial domain transcriptionally regulates *dyrk1b*, which further amplifies Fgf signaling by downregulating its feedback inhibitor, *spry1*, leading to specification of paraxial *myoD* domain (Fig.2K).

### Dyrk1b is not necessary but sufficient for adaxial *myoD* expression

The phenotypic similarity of cyclopic eyes between Dyrk1b-/- and the Sonic hedgehog (Shh)-/- embryos, and the essential role of Shh in specification of adaxial *myoD* domain (25, 49) led us to investigate if Dyrk1b interacts with Shh signaling to induce adaxial *myoD* expression. Additionally, DYRK1B has been shown to modulate switch between autocrine to paracrine Hedgehog signaling in pancreatic cancer cells (50). The adaxial cells are precursors of slow muscle fibers specified adjacent to the notochord and eventually migrate to the most lateral part of the somites (51). Since the adaxial *myoD* domain appeared intact in the Dyrk1b-/- embryos (Fig.1 X, Y), we investigated whether the sensitized background of Shh inhibition would reveal any novel interactions of Shh with Dyrk1b. We performed a series of experiments, starting with Shh inhibition using the well-established smoothened inhibitor cyclopamine (52). We treated the embryos at 10hpf to circumvent any effects of Shh inhibition during germ-layer formation in gastrulation. Cyclopamine treatment of embryos resulted in loss of adaxial *myoD* with only a small reduction of paraxial *myoD* (Fig. 2L, M) and recapitulated the *myoD* expression in smoothened mutants (25). To determine if Dyrk1b could rescue the adaxial *myoD* expression, we injected human *dyrk1b* mRNA at one-cell stage and treated with cyclopamine at 10hpf followed by examination of *myoD* at 10 somite stage. Strikingly, human *dyrk1b* mRNA considerably rescued the adaxial *myoD* (Fig.2N) (68.9% ± 1.37% were rescued, P-value =0.0003, average 40 ± 5 embryos, n=3 independent experiments). To explore the mechanisms by which Dyrk1b interacts with Shh signaling pathway, we first examined the expression of Shh ligand itself which revealed no significant difference in Shh expression in the notochord between Dyrk1b-/- and controls by WISH (Fig. 1A.4-A.5), suggesting that Dyrk1b does not affect the transcription of the Shh ligand. Likewise, the mRNA expression of Dyrk1b was unaffected upon cyclopamine treatment, suggesting that Shh signaling does not transcriptionally regulate Dyrk1b (data not shown). Next, we examined the effect of *dyrk1b* genetic manipulation on Shh effectors, Gli1, Gli2 and Gli3. In the Dyrk1b knockdown embryos at 14hpf, the protein levels of Gli1, and full length (active forms) Gli2 and Gli3 were significantly reduced as assayed by western blot (Number of independent experiments =2, technical replicates=2, pooled 50 embryos, Fig.2O, P). Similar patterns in Shh downstream effectors were obtained for Dyrk1b knockout embryos at 24hpf (SFig.3D). Conversely, the overexpression of Dyrk1b increased the protein levels of Gli1 at 14hpf (Number of independent experiments =2, technical replicates=2, pooled more than 50 embryos, Fig. 2O, P). These findings indicate that Dyrk1b rescued adaxial *myoD* in Shh-inhibited embryos by increasing the protein levels of Gli1. The suppression of Shh effectors in Dyrk1b knockout embryos, did not cause any substantial changes in the adaxial *myoD* domain leading to the conclusion that Dyrk1b is not necessary, but sufficient for adaxial *myoD* expression.

### The investigation of myocyte differentiation in C2C12 cells

In parallel with studies in zebrafish, we examined the interaction of Dyrk1b with Fgf and Shh signaling pathways in the mouse myoblast C2C12 cell line. C2C12 cells were previously isolated from 2-month old normal C3H mice, 70 hours post crush injury of the thigh muscle. The injury caused a rapid proliferation of myogenic mono-nucleated progenitors in the primary cultures from the muscle explants. These cells under low-serum conditions differentiated and fused into multi-nucleated myofibers within 3-4 days (53). These proliferating myogenic progenitors have been also described in the literature as “satellite cells” that can be stimulated to differentiate in low-serum conditions (54). Note that in zebrafish we examined the function of Dyrk1b in specification of myoblasts from pre-myogenic precursors while in C2C12 cells we are examining differentiation of committed myoblasts to myofibers. This may explain some of the key differences in the interaction of Dyrk1b with Fgf and Shh signaling pathways, as described below.

### The kinase activity of Dyrk1b is essential for differentiation of C2C12 myoblasts to myofibers

We transfected C2C12 cells with vectors containing GFP, *DYRK1B*^*WT*^, *DYRK1B*^*R102C*^ the most common human disease-causing mutation (10), and the kinase defective *DYRK1B*^*Y273F, K140R*^ at the beginning of differentiation and examined Myosin heavy chain 1 (MyHC1) expression at day 7 of differentiation. Dyrk1b levels were elevated in *DYRK1B*^*WT*^, *DYRK1B*^*R102C*^, *DYRK1B*^*Y273F, K140R*^ relative to GFP controls (SFig3B). A 2.69-fold higher percentage area of MyHC1 and 2.00 fold elevated nuclei per myofiber were observed in *DYRK1B*^*WT*^ compared to controls (Fig. 3A-E, n=3 independent experiments). In the *DYRK1B*^*Y273F, K140R*^ transfected C2C12 cells, the percentage area of MyHC1 and number of nuclei per myofiber were similar to the control cells (Fig.3 A, D, E). These data indicate that DYRK1B promotes differentiation of myoblasts to myofibers in a kinase-dependent fashion. *DYRK1B*^*R102C*^ transfected C2C12 cells, however, exhibited near complete abrogation of muscle differentiation as visualized by percent MyHC1 and nuclei per myofiber (Fig. 3C, E, n=3 independent experiments) and suggested loss of function as a potential mechanism for sarcopenia in *DYRK1B*^*R102C*^ mutation carriers. The more severe phenotype of *DYRK1B*^*R102C*^, which demonstrated only a 50% reduction in the kinase activity (44), is likely due to the kinase-independent effects of the mutation that has been previously shown in cultured HepG2 cells (10). Overall, these findings suggest a contribution of a kinase-independent function of Dyrk1b in regulation of myofiber differentiation. This was not further explored due to lack of available methods to target the kinase independent function for translational intervention.

**Figure 3:**
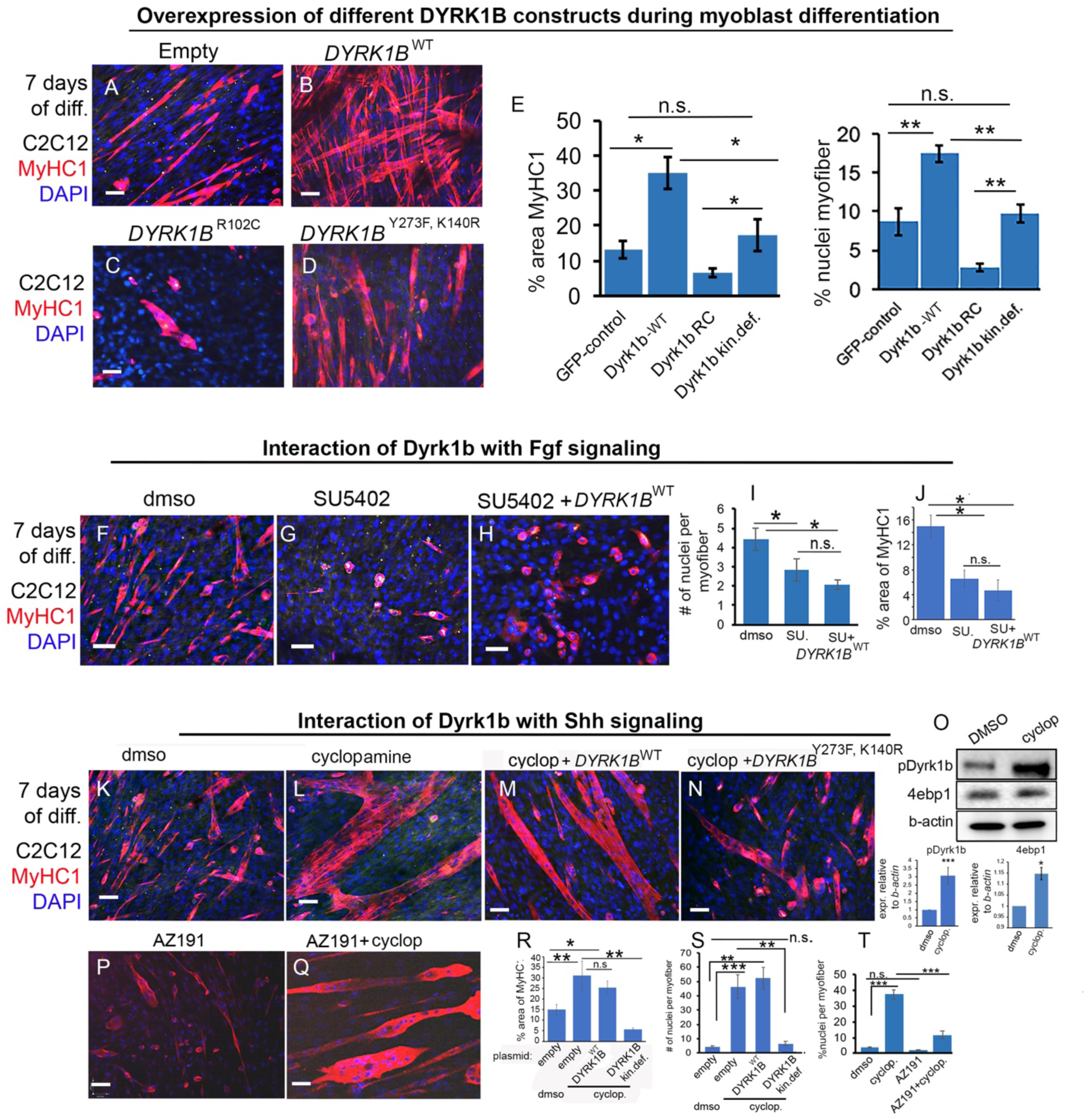
Interaction of Dyrk1b with Fgf and Shh signaling during specification of myofibers from myoblasts in C2C12 cells. **A-D** IF staining for myosin heavy chain (MyHC1) in C2C12 cells after 7 days of differentiation. The cells were transfected with plasmids containing no external cDNA but GFP, *DYRK1B*^*WT*^, *DYRK1B* ^*R102C*^ and *DYRK1B* ^*Y273F, K140R*^ (kinase-defective) before differentiation was ensued. **E** Quantification for % area of MyHC1 (mean ± s.e.m) and nuclei per myofiber (mean ± s.e.m) on 7^th^ day of differentiation in the indicated conditions. **F-H** IF staining of MyHC1 expression on 7th day of C2C12 differentiation in cells treated with dmso (H), SU5402 (I) and SU5402 + *DYRK1B*^WT^. **I**. Quantification of the nuclei (mean ± s.e.m) per myofiber in the indicated conditions. **J** Percentage area (mean ± s.e.m) of MyHC1 expression on 7^th^ day of differentiation. The cells were transfected before beginning the differentiation and treated with low dose of SU5402 for all seven days of differentiation. **K-N** IF staining of MyHC1 at 7th day of differentiation in C2C12 cells in dmso control (K), cyclopamine transfected with empty vector (no cDNA) (L); cyclopamine and wild type *DYRK1B* (M) or cyclopamine and kinase defective *DYRK1B* (N). **O.** Western blot and quantification of phosphorylated DYRK1B and 4e-bp1 in cyclopamine treated C2C12 cells. **P-Q**, IF staining of MyHC1 in C2C12 cells treated with inhibitor AZ191 alone or co-treated with inhibitor AZ191 and cyclopamine. **R-S** % area (mean ± s.e.m) of MyHC1(R) and quantification of % nuclei (mean ± s.e.m) of myofibers for the panels K-N **T.** quantification of % nuclei (mean ± s.e.m) per myofibers for the panels P, Q. * indicates p ≤0.05, ** p≤ 0.01, *** p≤ 0.001. Students ttest, 2 tailed. Scale bar = 200µm.

### Exclusion of Dyrk1b as an effector of Fgf signaling in myofiber differentiation

We then investigated the interaction between Fgf and Dyrk1b during muscle differentiation in C2C12 myoblasts. We inhibited FGF signaling from day1 to day 7 of differentiation, after which cells were fixed for IF staining (number of independent experiments=3). To prevent toxicity, we used 1µM of SU5402, a substantially lower dose compared to those used in previous studies (55). This dose of SU5402 was sufficient to suppress myoblast differentiation as assessed by percentage of MyHC1 positive cells and myotubule fusion evident by nuclei per myofiber (Fig. 3F, G, I, J). The overexpression of *DYRK1B*^*WT*^ was unable to rescue loss of MyHC1 and myoblasts fusion (Fig. 3 H, I, J, n=3 independent experiments) in SU5402 treated cells. This finding suggests that either 1. other potent downstream effectors, in addition to Dyrk1b, mediate the effects of Fgf signaling during C2C12 myogenesis 2. Fgf and Dyrk1b independently regulate myoblast differentiation; with Fgf being a stronger stimulator of myogenesis.

### Dyrk1b is activated by Shh inhibition and promotes myotube fusion

The relationship between Dyrk1b and Shh in the context of myoblast fusion and differentiation is not known. Both inhibitory and stimulating effects of Shh activation on myoblast fusion have been suggested (27, 35), which is likely due to different experimental conditions. We treated C2C12 myoblasts with cyclopamine at a dose of 1µM for all 7 days of differentiation. This cyclopamine treatment of C2C12 myoblasts resulted in increased MyHC1 levels and excessive myoblast fusion with the number of nuclei per fiber increasing by about 10-fold on day7 of differentiation compared to vehicle alone (DMSO= 4.42 ± 0.6 nuclei, Cyclopamine= 46.44± 8.2 nuclei per fiber, respectively, p= 9.0×10^−4^, number of independent experiments=3, Fig. 3 K,L, R, S). To determine if myoblast fusion induced by cyclopamine is dependent on kinase activity of Dyrk1b, we overexpressed control vector, *DYRK1B*^*WT*^, and *DYRK1B*^*Y273F, K140R*^ in C2C12 cells along with cyclopamine. The excess myoblast fusion induced by cyclopamine was blocked in C2C12 cells expressing *DYRK1B*^*Y273F, K140R*^ but not with *DYRK1B*^*WT*^ (Fig. 3 M, N, R, S). To further confirm the role of Dyrk1b kinase activity in myoblast fusion, we co-treated C2C12 cells with cyclopamine and AZ191, a specific kinase inhibitor of Dyrk1b (56) for all 7 days of differentiation. We tested several concentrations of AZ191 and found 800nM to be the lowest concentration that reduces pDyrk1b levels (SFig.3E). Co-treatment of cyclopamine with AZ191 reduced the number of nuclei per myofiber by 75% compared to cyclopamine alone (Fig. 3 P, Q, T). Altogether, these findings suggest that Shh inhibition increased myotube fusion in a manner dependent on the kinase function of Dyrk1b. Next, we hypothesized that increased auto-phosphorylation of Dyrk1b via Shh-inhibition is likely responsible for its enhanced catalytic activity since the formation of catalytic active Dyrk1b is dependent on its auto-phosphorylation (57). We examined phospho-Dyrk1b (p-Y273) levels in C2C12 cells after treatment with cyclopamine. The antibody against p-HIPK2 cross-reacts with the phospho-Y273 residue of the Dyrk1b protein and was previously used to assess Dyrk1b activation (45). WB analysis in cyclopamine treated C2C12 cells revealed higher Dyrk1b auto-phosphorylation upon Shh inhibition compared to untreated cells (Fig. 3O). Collectively, our findings indicate that myofiber differentiation of C2C12 cells requires auto-phosphorylation of Dyrk1b, which can be achieved by inhibiting Shh signaling.

### The proteomics analysis identifies the augmented muscle differentiation pathways in Dyrk1b overexpressing C2C12 cells

The downstream effectors of Dyrk1b during skeletal muscle differentiation are largely unknown. To this end, we carried out proteomic analysis of C2C12 cells in DYRK1B^WT^ and the corresponding controls. The expression of 362 proteins was significantly altered by DYRK1B overexpression in C2C12 cells compared to the controls (Table1, Fig. 4 A, B, n=3 biological replicates, unique peptides >2, ttest ≤0.05, FDR ≤0.05, Benjamini-Hochberg correction, fold change ≤0.5 and ≥1.5). The Principal Component Analysis showed segregation of the 6 samples into 2 distinct groups: one group for the 3 controls and the other for 3 Dyrk1b overexpression samples indicating that these two sets of samples are distinct in their expression patterns. Ingenuity Pathway Analysis was used to decipher the changes in the pathways upon Dyrk1b overexpression in C2C12 cells. The top canonical pathways with p-value of <0.05 and z-score of ≤-2 or ≥ 2, included activation of Glycolysis, AMPK Signaling, Rho-mediated Signaling, Actin Cytoskeletal Signaling and Integrin Signaling in DYRK1B^WT^ (Table2, Fig.4C). Prominent proteins most relevant to muscle differentiation in these pathways included Pard3 (elevated 1897 fold), SRC non-receptor kinase (elevated 8.89), Focal Adhesion Protein1 (elevated 7.87 fold), Arhgef17 (elevated 5.9 fold), Protein Phosphatase 1B (reduced 1.85), Cofilin1 (reduced 1.14), and elevation of several isoforms of myosin: Myosin Light chain 1/3 (slow twitch, elevated 2.32) and Troponin T (slow twitch, elevated 4.42), Troponin T (fast twitch fiber, elevated 1.77) (Table1). These changes in the myosin and troponin isoforms are consistent with microarray analysis previously reported in C2C12 cells when Dyrk1b was knocked down (13). IPA additionally found Egf, Vegf and chemokine signaling, which have been implicated in increasing growth and metabolic capacity of the myocytes (39, 58), to be activated in DYRK1B^WT^ cells. Further, skeletal muscle contractility, cell movements and glycolysis were predicted to be increased and myofiber degeneration was predicted to be decreased (Fig.4D, Table3) in DYRK1B^WT^ cells. The proteomic analysis also predicted the potential upstream regulators of the proteins that were significantly altered (Fig.4E). Strikingly, MyoD (z-score 2.71) and Mef2c (z-score 2.09), the two important transcription factors for muscle differentiation, were identified to be activated in DYRK1B^WT^ C2C12 cells (Table4, Fig.4E). Remarkably, p38 MAPK (z-score 2.77) (59), EgfR (z-score 2.48) (60) and Insulin (2.01), previously implicated in muscle differentiation, were also predicted to be activated (Table4). Altogether, a complex set of several regulators of muscle differentiation were altered upon elevated Dyrk1b expression in C2C12 cells.

**Table 1:**
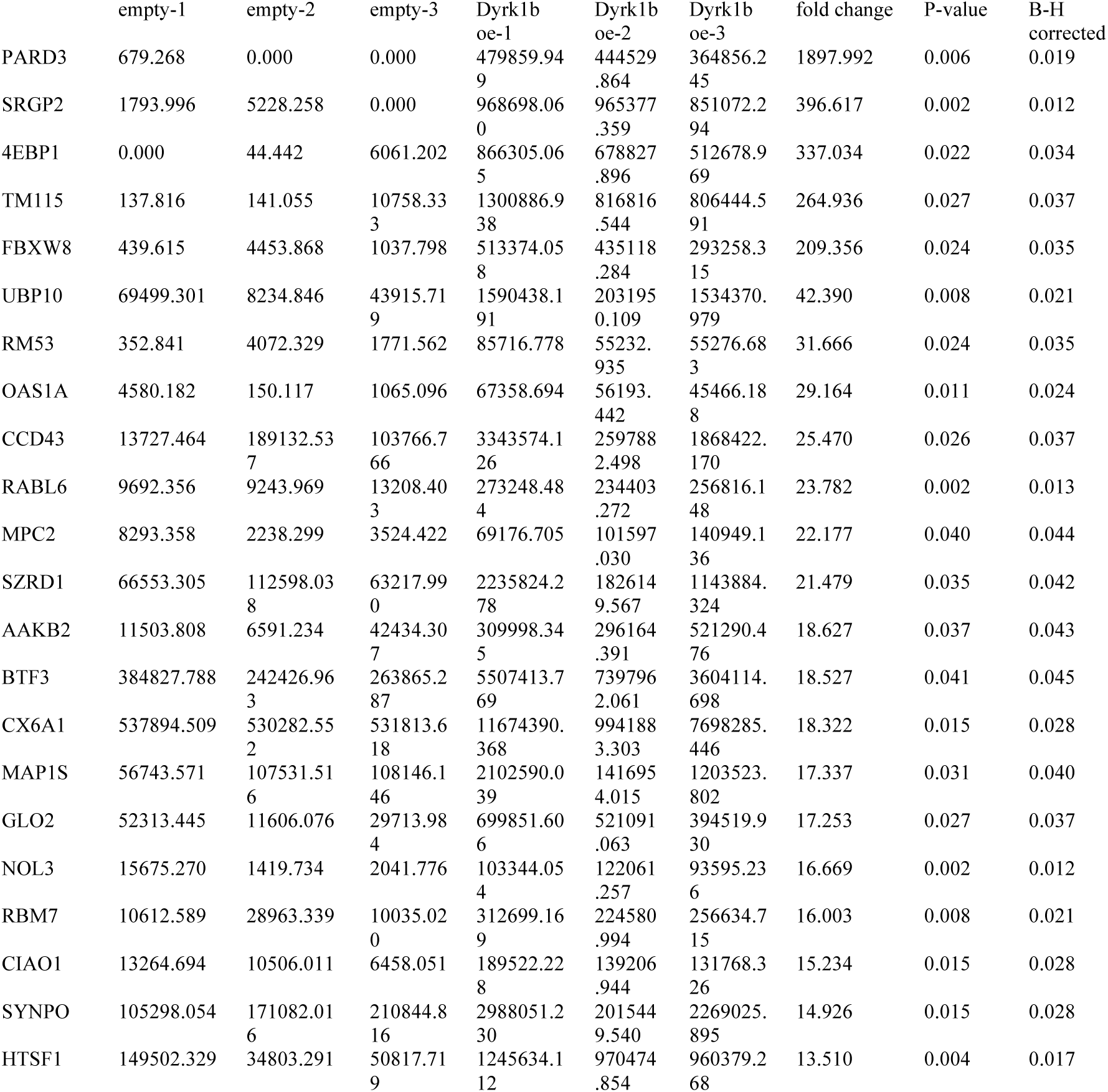
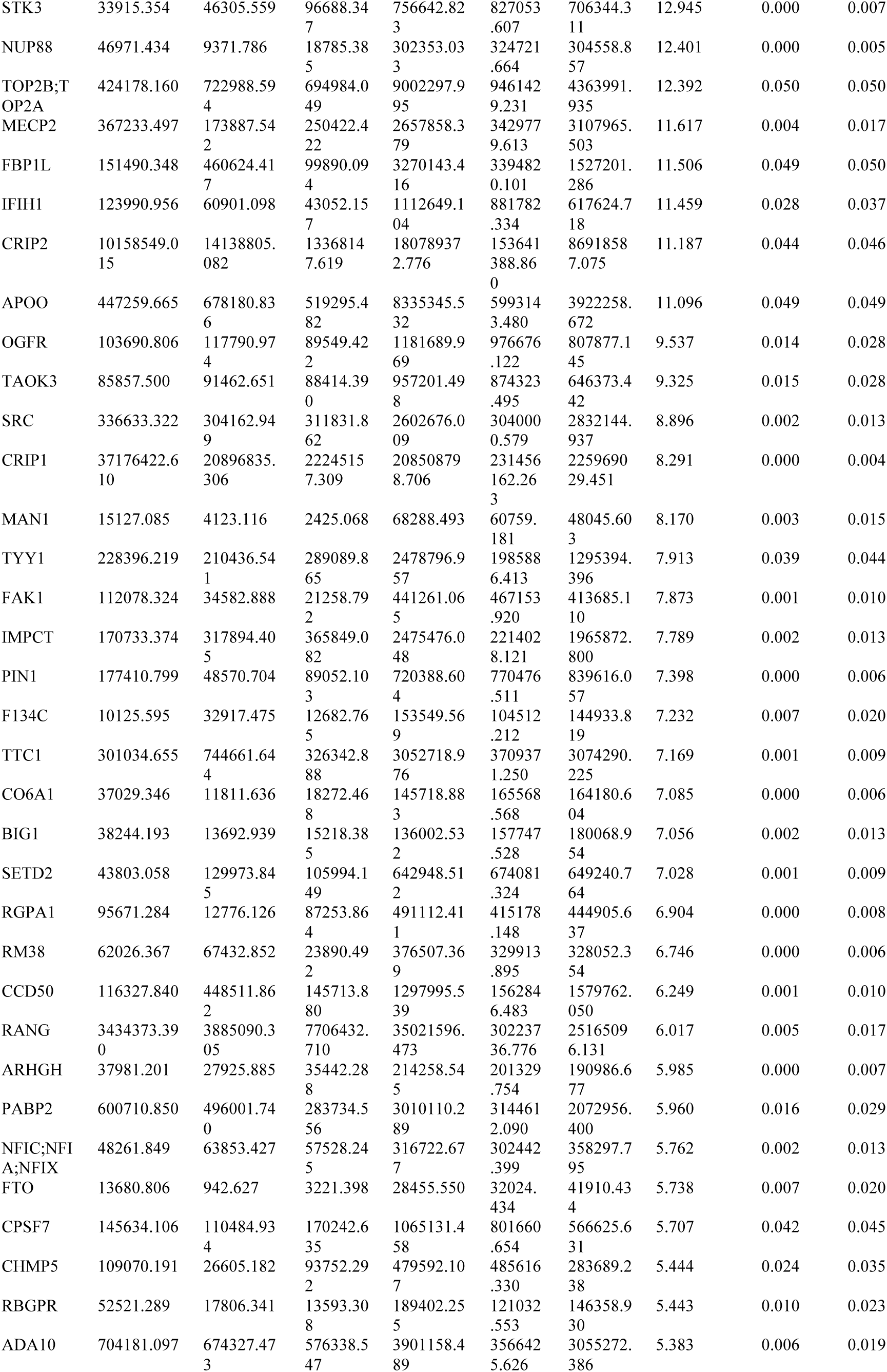
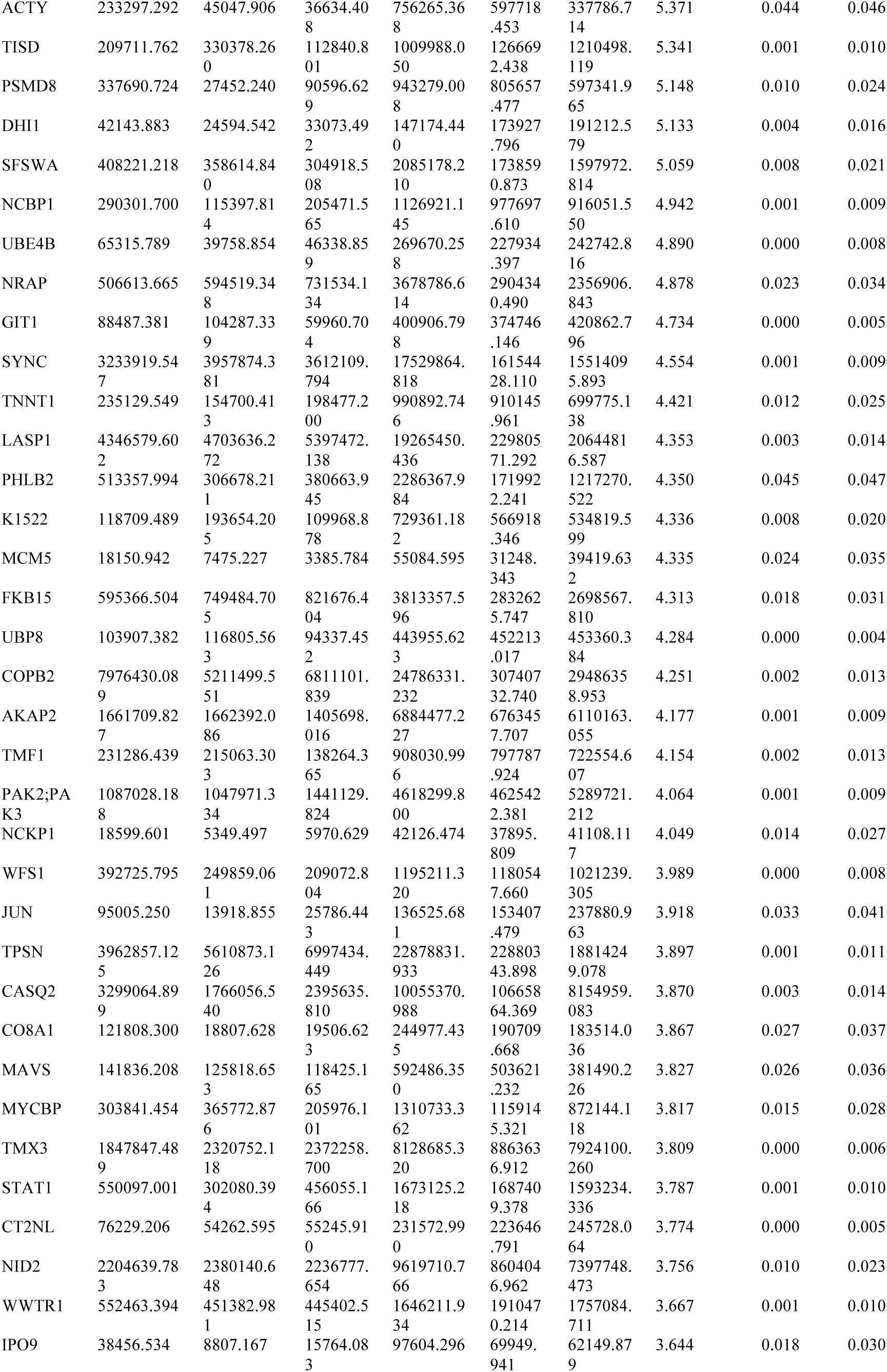
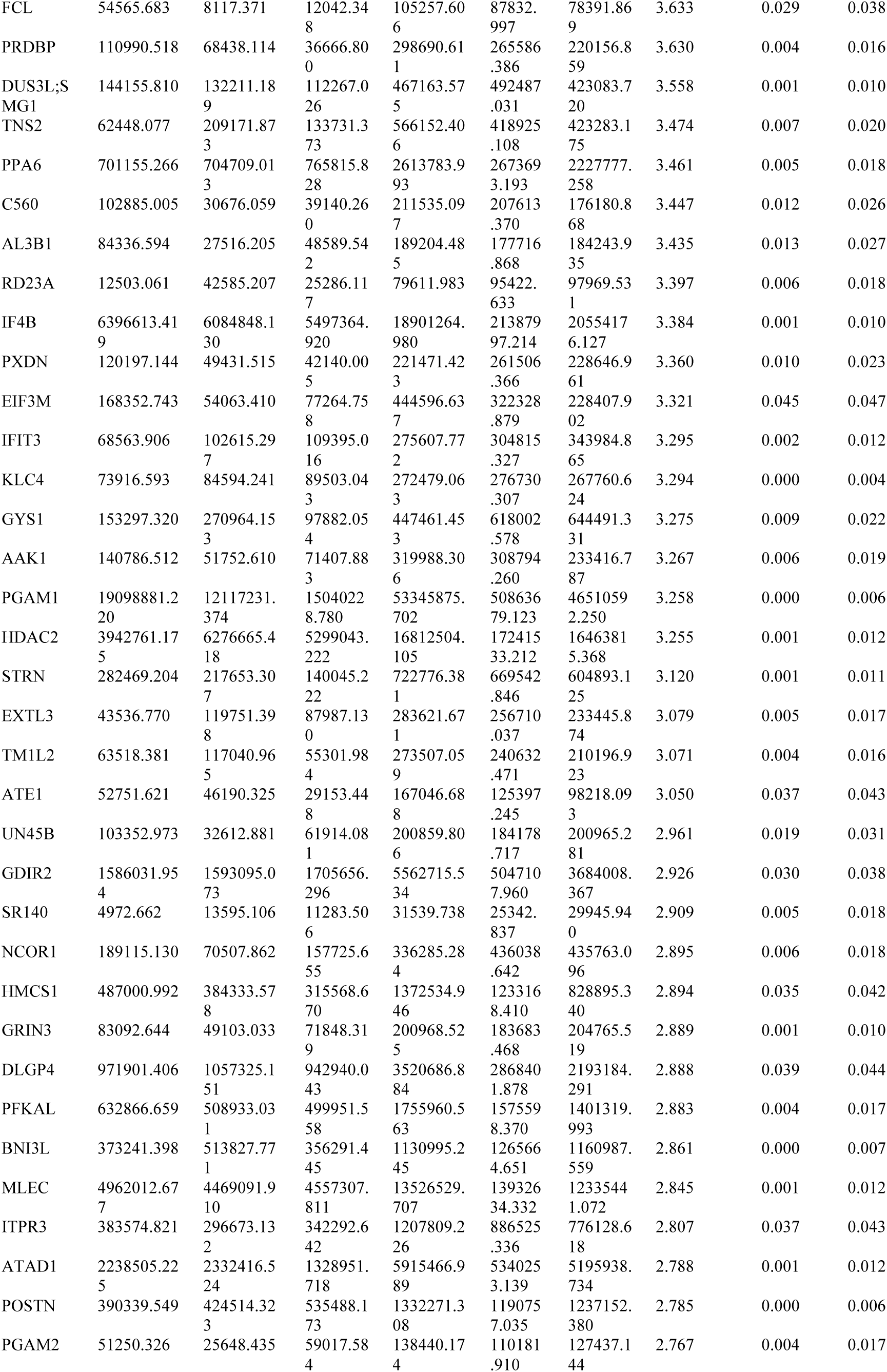
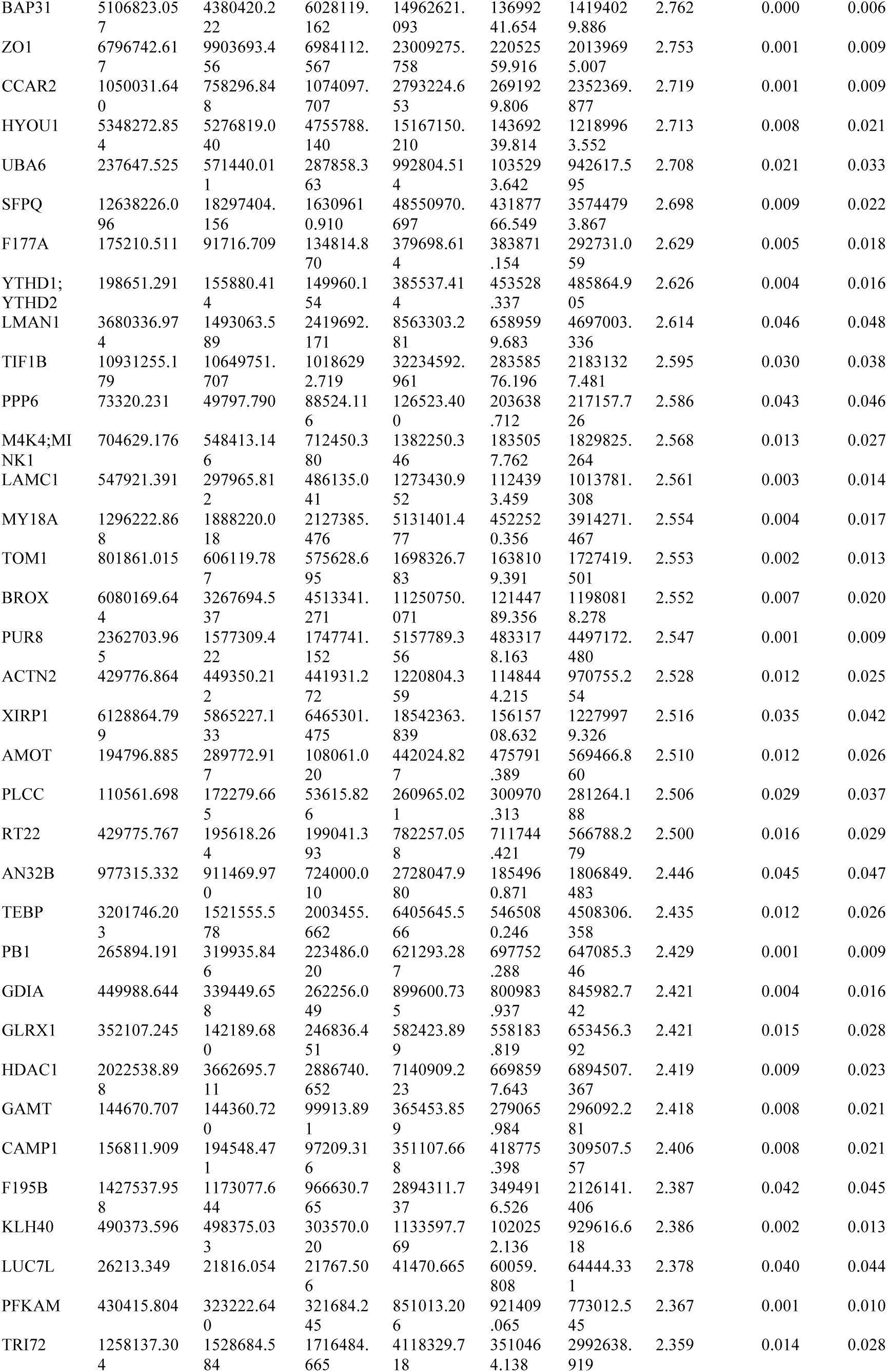
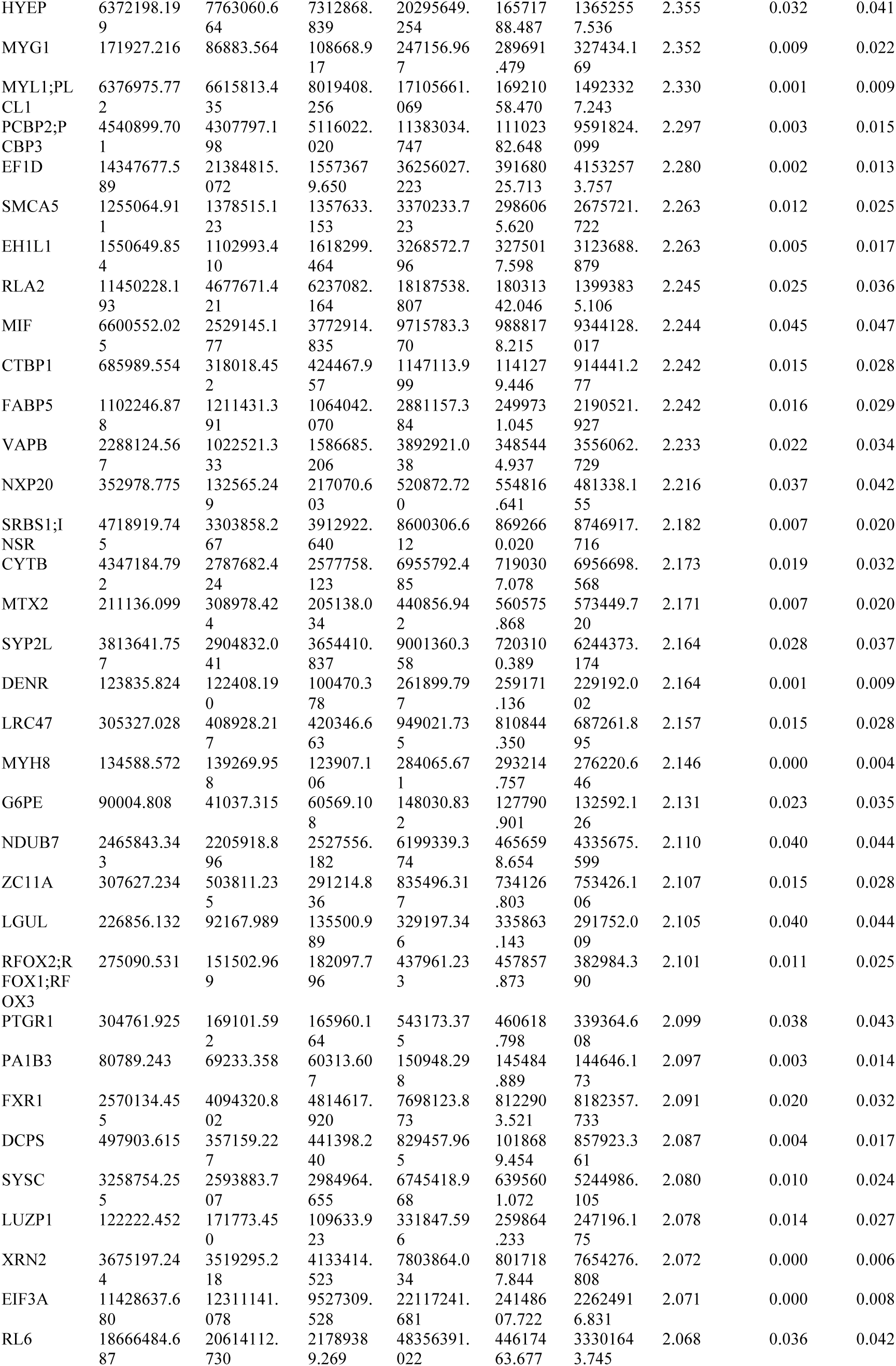
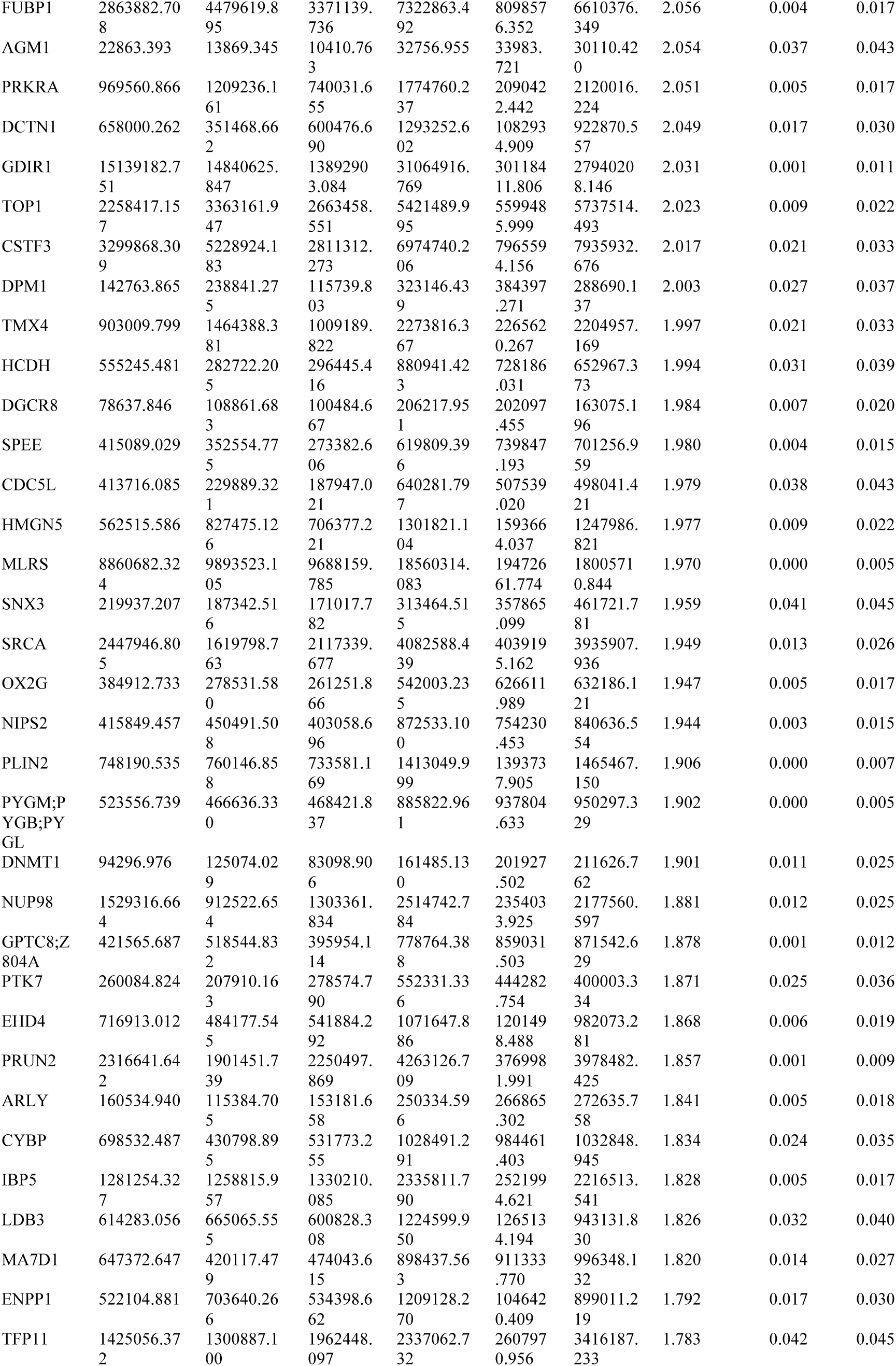
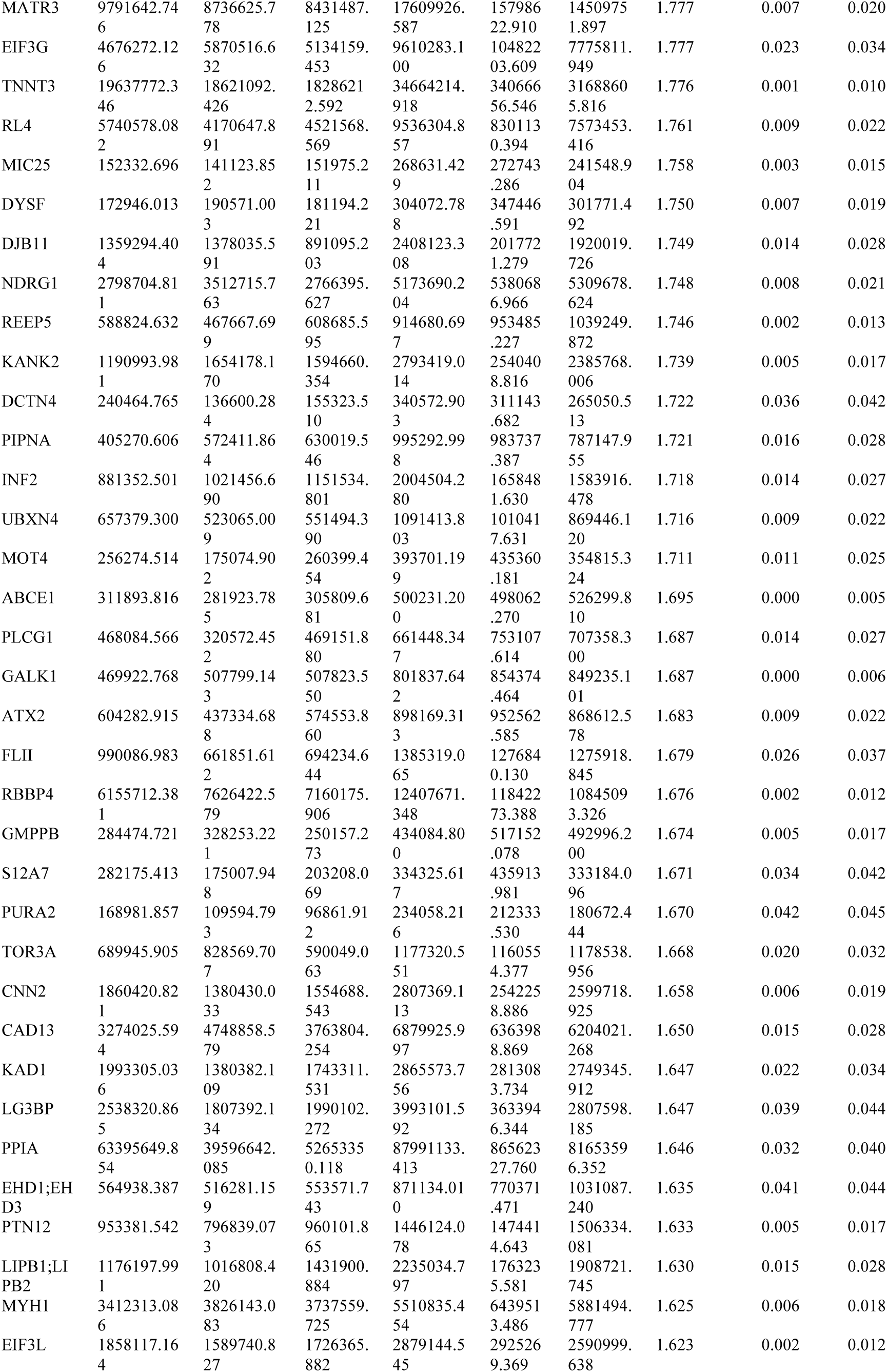
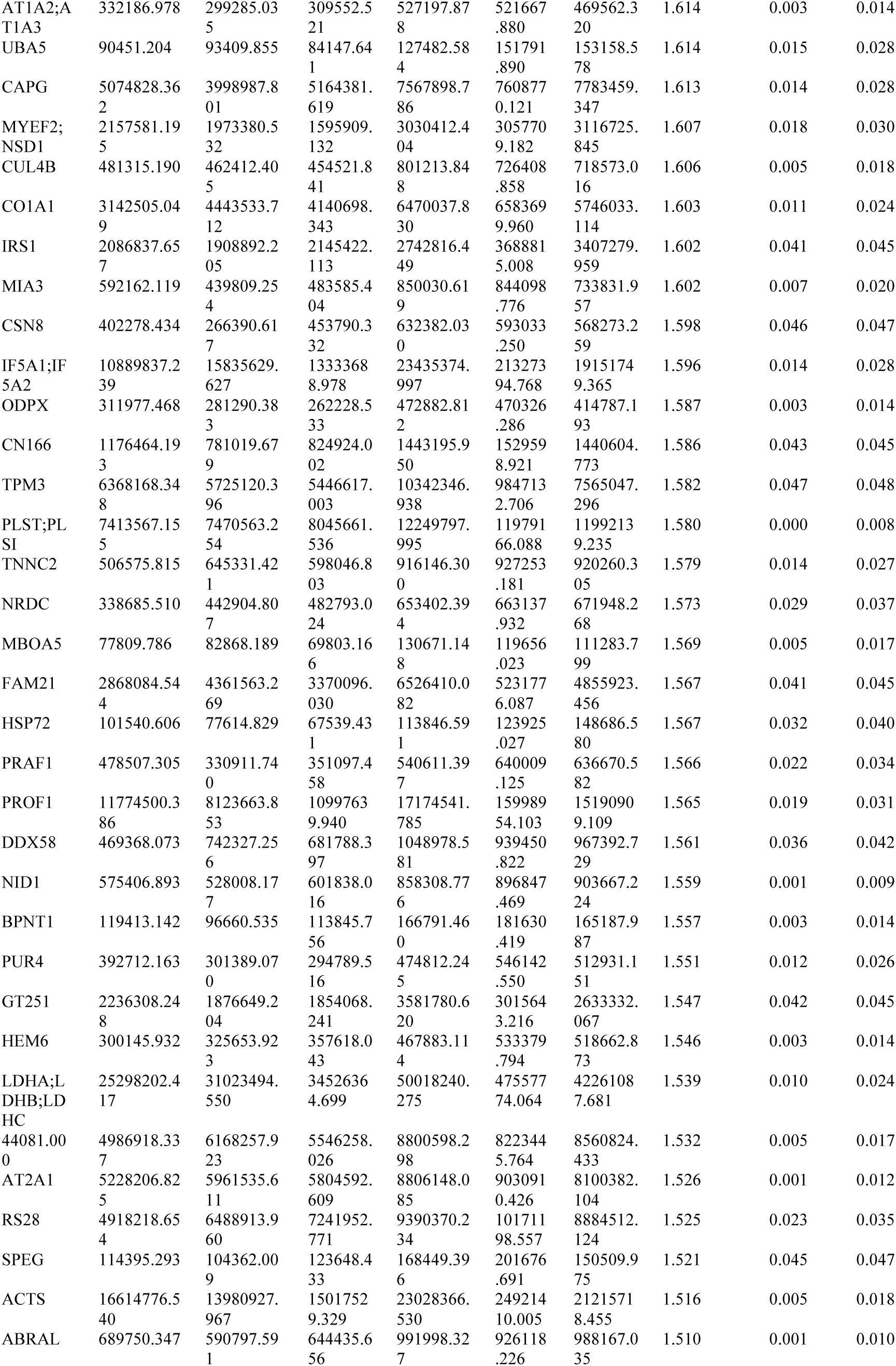
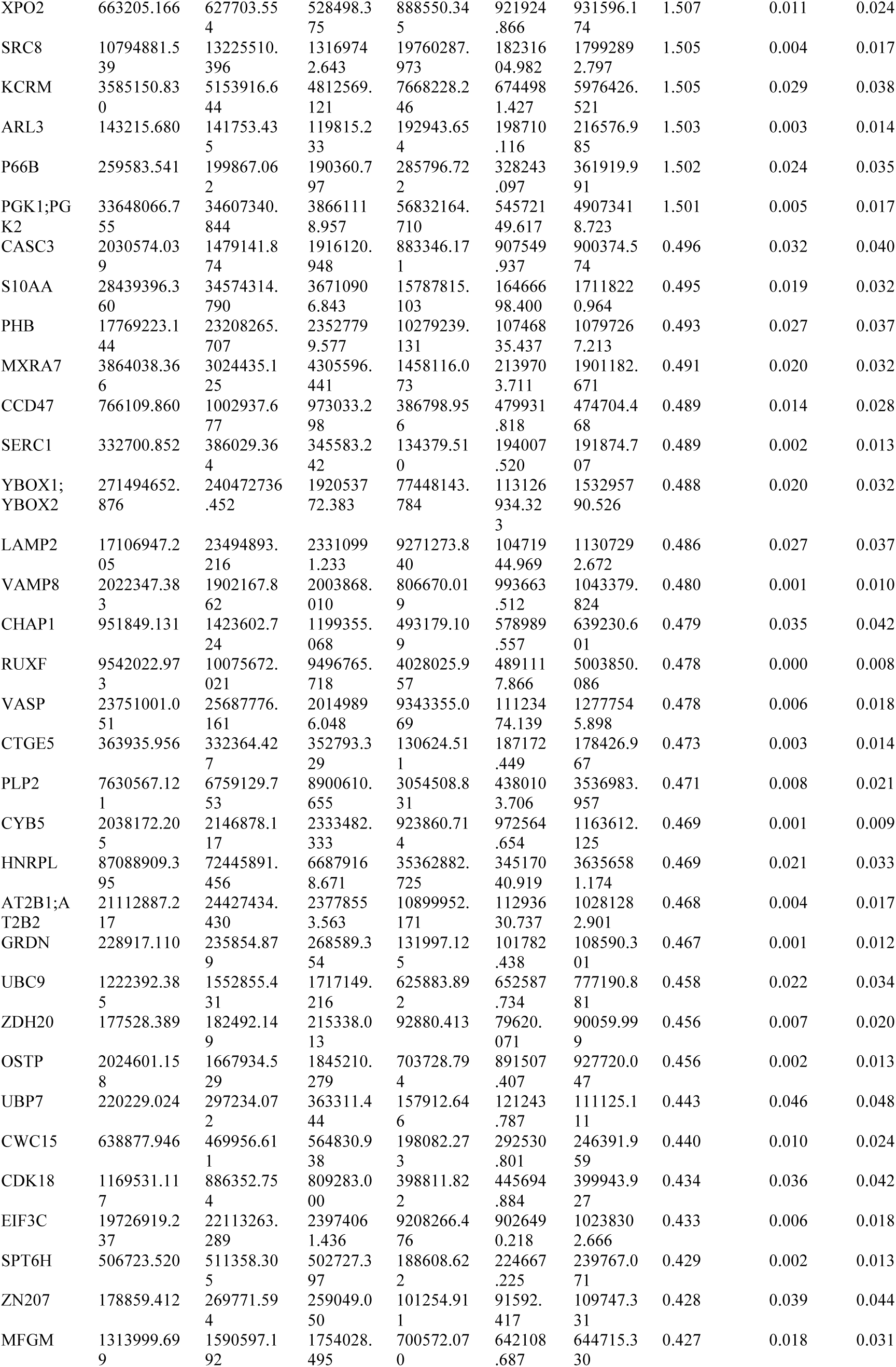
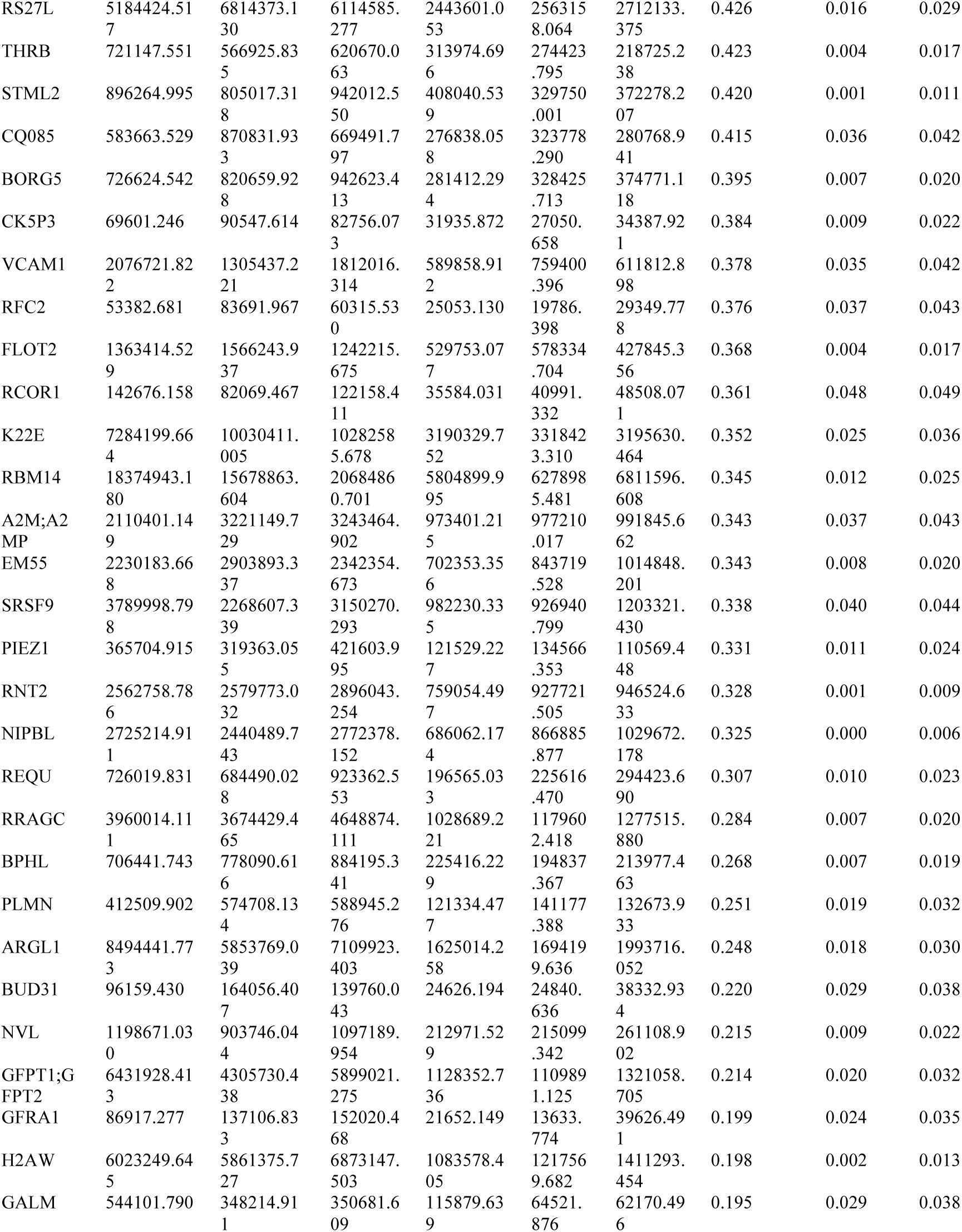
The list of significantly proteins altered in C2C12 cells upon Dyrk1b overexpression. The cells were serum starved on day 7 of differentiation prior to subjecting them to proteomics by LC-MS/MS.

**Table 2:**
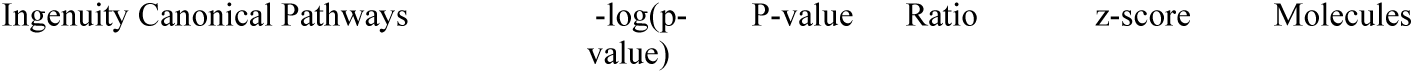
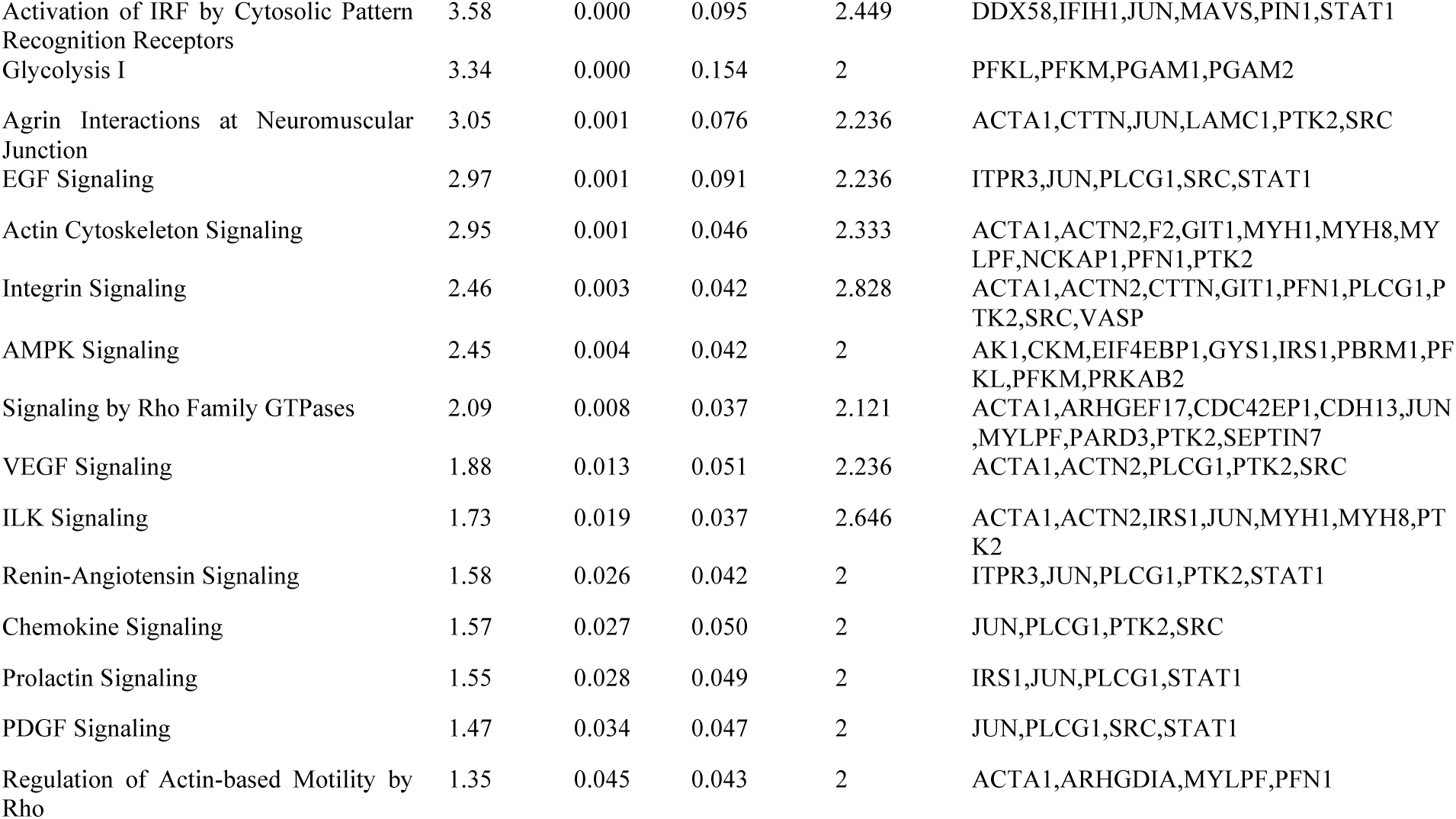
The top canonical pathways predicted by Ingenuity Pathway Analysis. The significantly altered proteins (FDR ≤0.05, fold change≤ 0.05, ≥1.5) were used as inputs. The Pathways were then filtered again by their p-values and z-scores of >2, <-2.

|  |  |  |  |  |  |
| --- | --- | --- | --- | --- | --- |
| Activation of IRF by Cytosolic Pattern Recognition Receptors | 3.58 | 0.000 | 0.095 | 2.449 | DDX58,IFIH1,JUN,MAVS,PIN1,STAT1 |
| Glycolysis I | 3.34 | 0.000 | 0.154 | 2 | PFKL,PFKM,PGAM1,PGAM2 |
| Agrin Interactions at Neuromuscular Junction | 3.05 | 0.001 | 0.076 | 2.236 | ACTA1,CTTN,JUN,LAMC1,PTK2,SRC |
| EGF Signaling | 2.97 | 0.001 | 0.091 | 2.236 | ITPR3,JUN,PLCG1,SRC,STAT1 |
| Actin Cytoskeleton Signaling | 2.95 | 0.001 | 0.046 | 2.333 | ACTA1,ACTN2,F2,GIT1,MYH1,MYH8,MYLPF,NCKAP1,PFN1,PTK2 |
| Integrin Signaling | 2.46 | 0.003 | 0.042 | 2.828 | ACTA1,ACTN2,CTTN,GIT1,PFN1,PLCG1,PTK2,SRC,VASP |
| AMPK Signaling | 2.45 | 0.004 | 0.042 | 2 | AK1,CKM,EIF4EBP1,GYS1,IRS1,PBRM1,PFKL,PFKM,PRKAB2 |
| Signaling by Rho Family GTPases | 2.09 | 0.008 | 0.037 | 2.121 | ACTA1,ARHGEF17,CDC42EP1,CDH13,JUN,MYLPF,PARD3,PTK2,SEPTIN7 |
| VEGF Signaling | 1.88 | 0.013 | 0.051 | 2.236 | ACTA1,ACTN2,PLCG1,PTK2,SRC |
| ILK Signaling | 1.73 | 0.019 | 0.037 | 2.646 | ACTA1,ACTN2,IRS1,JUN,MYH1,MYH8,PTK2 |
| Renin-Angiotensin Signaling | 1.58 | 0.026 | 0.042 | 2 | ITPR3,JUN,PLCG1,PTK2,STAT1 |
| Chemokine Signaling | 1.57 | 0.027 | 0.050 | 2 | JUN,PLCG1,PTK2,SRC |
| Prolactin Signaling | 1.55 | 0.028 | 0.049 | 2 | IRS1,JUN,PLCG1,STAT1 |
| PDGF Signaling | 1.47 | 0.034 | 0.047 | 2 | JUN,PLCG1,SRC,STAT1 |
| Regulation of Actin-based Motility by Rho | 1.35 | 0.045 | 0.043 | 2 | ACTA1,ARHGDIA,MYLPF,PFN1 |

**Table 3:** The cellular and molecular functions altered by Dyrk1b overexpression in C2C12 cells. The different categories were filtered by their p-values ≤0.05 and z-scores of >2, <-2.

| Diseases or Functions Annotation | p-value | Predicted Activation State | Activation z-score |
| --- | --- | --- | --- |
| Size of body | 0.00000 | Increased | 4.511 |
| Infection by HIV-1 | 0.00101 | Increased | 4.087 |
| Infection of cells | 0.00002 | Increased | 4.068 |
| Infection by RNA virus | 0.00057 | Increased | 3.94 |
| Quantity of cells | 0.00002 | Increased | 3.442 |
| Infection of epithelial cell lines | 0.00009 | Increased | 3.299 |
| Infection of kidney cell lines | 0.00014 | Increased | 3.299 |
| Infection of embryonic cell lines | 0.00014 | Increased | 3.299 |
| Viral Infection | 0.00000 | Increased | 3.176 |
| Cell survival | 0.00000 | Increased | 2.674 |
| Contractility of skeletal muscle | 0.00000 | Increased | 2.414 |
| Cell movement of epithelial cell lines | 0.00041 | Increased | 2.37 |
| Cell viability | 0.00001 | Increased | 2.293 |
| Cell movement of stem cells | 0.00131 | Increased | 2.226 |
| Antiviral response of cells | 0.00063 | Increased | 2.219 |
| Biosynthesis of purine ribonucleotide | 0.00109 | Increased | 2.219 |
| Hydrolysis of ATP | 0.00075 | Increased | 2.213 |
| Contractility of muscle | 0.00000 | Increased | 2.21 |
| Glycolysis of cells | 0.00092 | Increased | 2.2 |
| Migration of epithelial cell lines | 0.00009 | Increased | 2.151 |
| Cell cycle progression | 0.00001 | Increased | 2.143 |
| Quantity of blood cells | 0.00078 | Increased | 2.139 |
| Interaction of protein | 0.00035 | Increased | 2.095 |
| Invasion of breast cancer cell lines | 0.00013 | Increased | 2.084 |
| Metabolism of nucleic acid component or derivative | 0.00097 | Increased | 2.032 |
| Metabolism of nucleotide | 0.00161 | Increased | 2.032 |
| Degeneration of myofiber | 0.00010 | Decreased | -2 |
| Cell death of cancer cells | 0.00126 | Decreased | -2.04 |
| Necrosis | 0.00000 | Decreased | -2.086 |
| Necrosis of muscle | 0.00046 | Decreased | -2.099 |
| Cell death of osteosarcoma cells | 0.00065 | Decreased | -2.121 |
| Degeneration of muscle cells | 0.00012 | Decreased | -2.219 |
| Movement Disorders | 0.00022 | Decreased | -2.368 |
| Cell death of tumor cells | 0.00012 | Decreased | -2.401 |
| Motor dysfunction or movement disorder | 0.00008 | Decreased | -2.73 |
| Growth Failure | 0.00032 | Decreased | -4.333 |
| Organismal death | 0.00000 | Decreased | -6.008 |

**Table 4:**
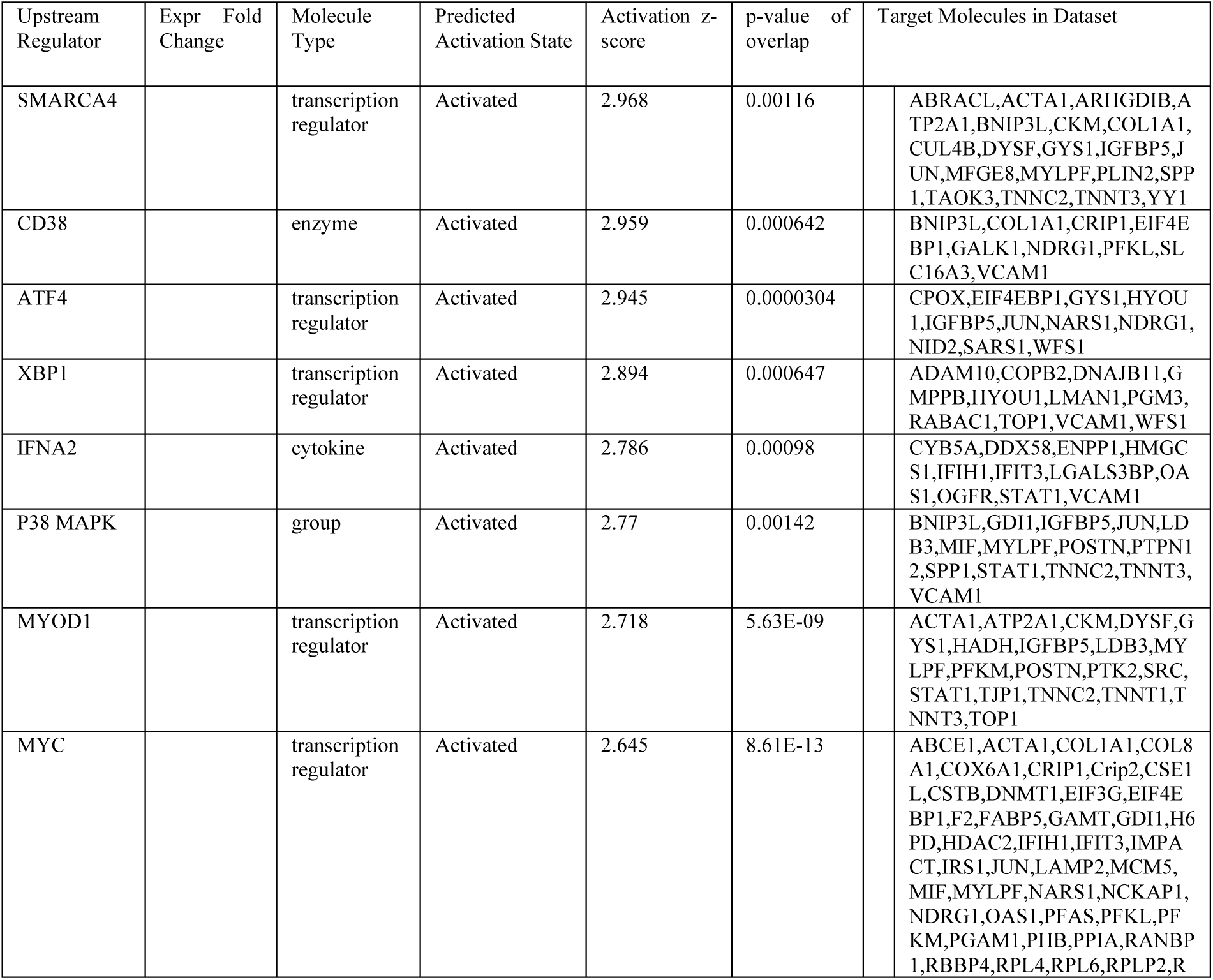
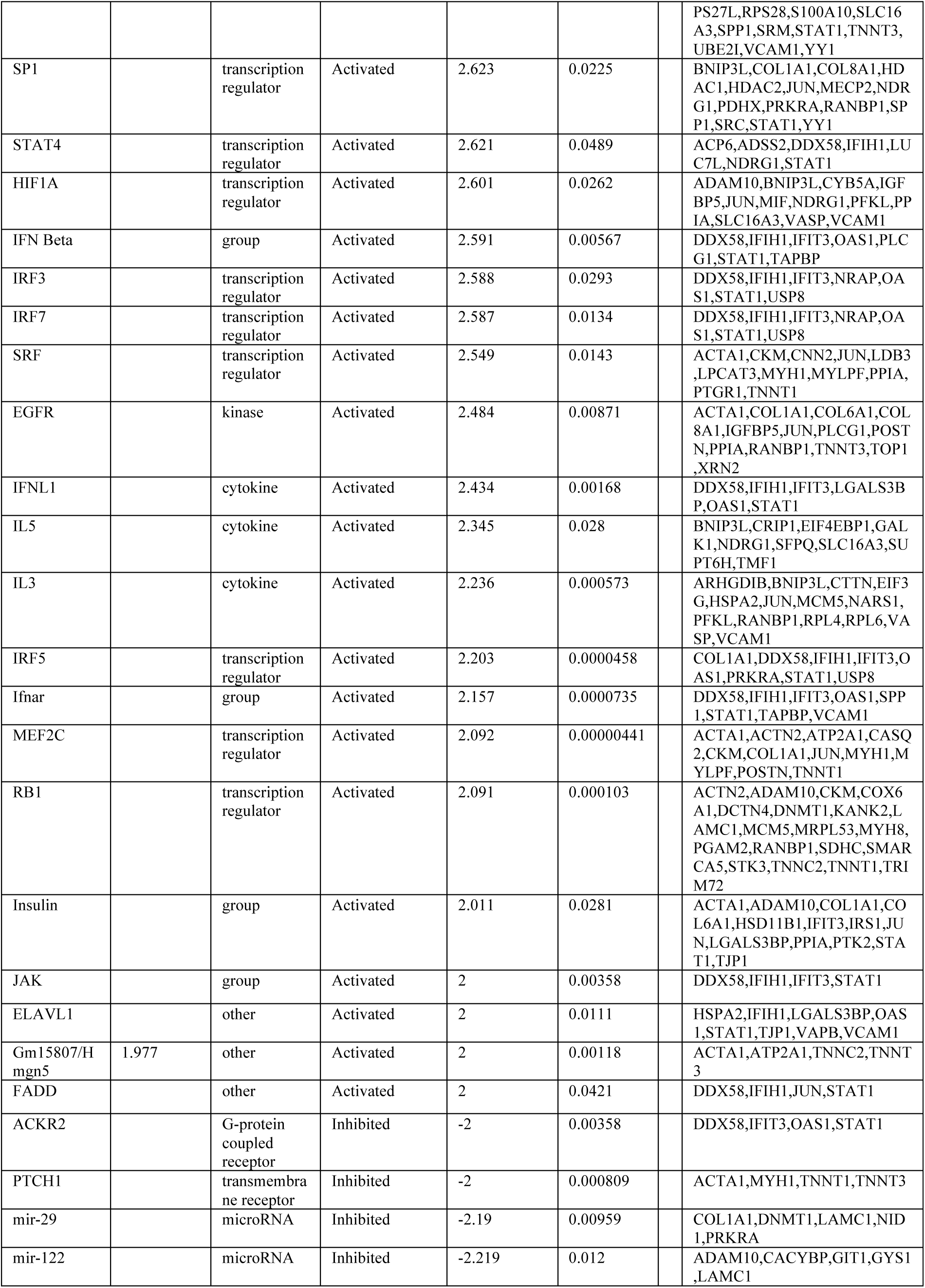
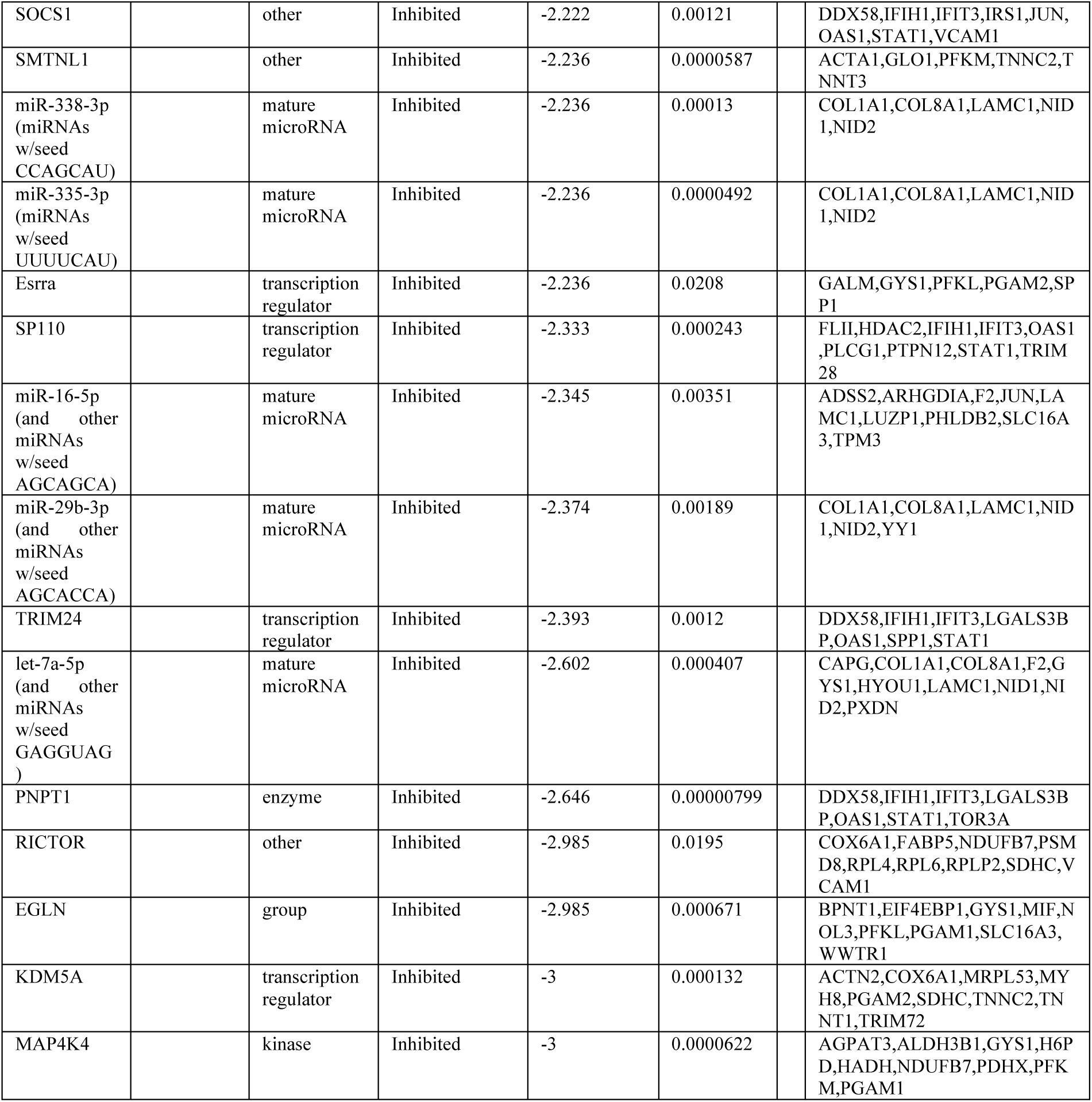
The predicted upstream regulators of proteins altered by Dyrk1b overexpression in C2C12 cells. The potential candidates were sorted by p-values ≤0.05 and z-scores of >2, <-2.

**Figure 4:**
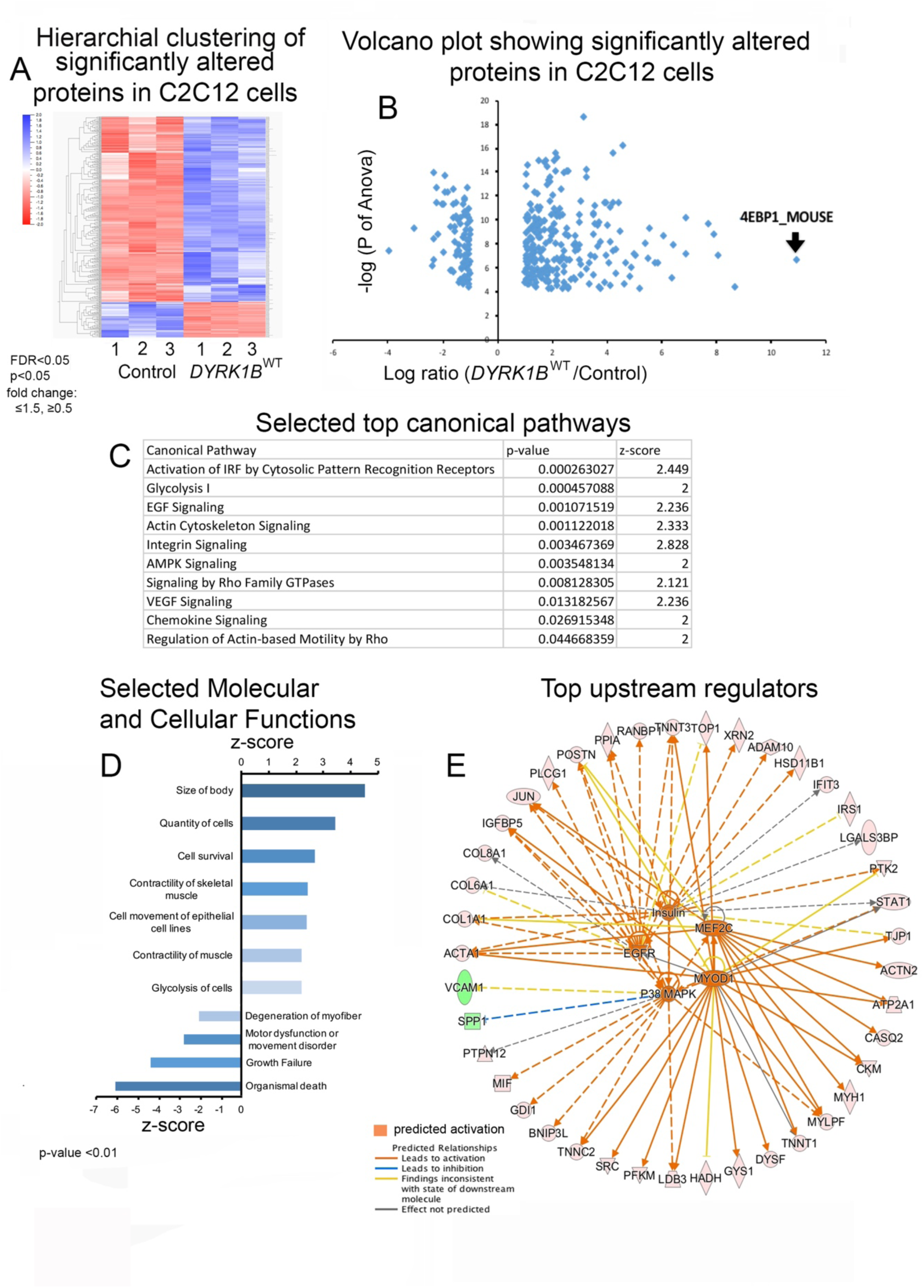
Proteomic analysis of C2C12 cells overexpressing Dyrk1b. **A** The significantly altered proteins in empty-vector transfected (N=3 biological replicates) and DYRK1B^WT^ transfected C2C12 cells (N=3 biological replicates) were hierarchically clustered using Qlucore-omics software. The significantly altered proteins were sorted by following thresholds: p≤0.05, false discovery rate (FDR) ≤0.05, Benjamini-Hochenberg corrected, fold change ≤0.5 or ≥1.5 fold. The cells were serum-starved prior to proteomics analysis. The coloring legend is besides the heat-map. **B** Volcano plot showing the spread of highly significantly altered proteins displayed in panel A (fold change < or >10, Anova P< 10^−4^. **C, D** All the significantly altered proteins (from panel A) were input into Ingenuity Pathway Analysis (IPA) and the top significantly altered pathways (C) and functions (D), with the z-scores (≥2 or ≤-2) are shown. **E** The most relevant selected upstream regulators, as predicted by IPA, of the significantly altered proteins are displayed. The orange color denotes activation of the molecule.

### Dyrk1b inhibits mTORC1, upregulates 4e-bp1 and activates autophagy

One of the most striking findings of the proteomics analysis was more than 300-fold increase in protein levels of eukaryotic translation initiation factor 4E (EIF4E), binding protein 1 (4e-bp1) in DYRK1B^WT^ vs. GFP transfected C2C12 cells (Fig.4A, B, Table1). Under normal conditions, 4e-bp1 inhibits the cap-dependent translation of mRNAs unless phosphorylated by mTORC1 complex (61). Paradoxically, elevated 4e-bp1 activity in the mouse skeletal muscle leads to increased MyHC1 expression and protects against high-fat induced Insulin resistance (62). We confirmed by WB that 4e-bp1 levels were elevated in C2C12 cells expressing Dyrk1b^WT^ and Dyrk1b^R102C^ relative to GFP-expressing controls on day 7 of differentiation (Fig. 5A, C, n=3 biological replicates). No difference was observed in *4e-bp1* mRNA levels between C2C12 cells overexpressing DYRK1B-WT vs. GFP-vector (data not shown), suggesting that Dyrk1b post-transcriptionally regulates 4e-bp1. We then examined whether 4ebp1 mediates the effects of Dyrk1b in promoting muscle differentiation in C2C12 cells. Indeed, the knockdown of 4e-bp1 using siRNA (Fig.5D) at the onset of differentiation prevented the increased myotubule formation in Dyrk1b^WT^ C2C12 cells (Fig. 5E-H, n=3 independent experiments). We further confirmed our findings *in-vivo*, where a complete loss of total 4e-bp1 protein (Fig. 5B, C) levels was observed in 24hpf Dyrk1b Crispr/Cas9 knockout zebrafish embryos (n=50 pooled embryos for WB). The overall protein synthesis was not previously reported to be affected when 4e-bp1 was overexpressed in the mouse skeletal muscle (62, 63). Altogether, these series of experiments indicate Dyrk1b increases myofiber specification by increasing 4e-bp1 protein expression.

**Figure 5:**
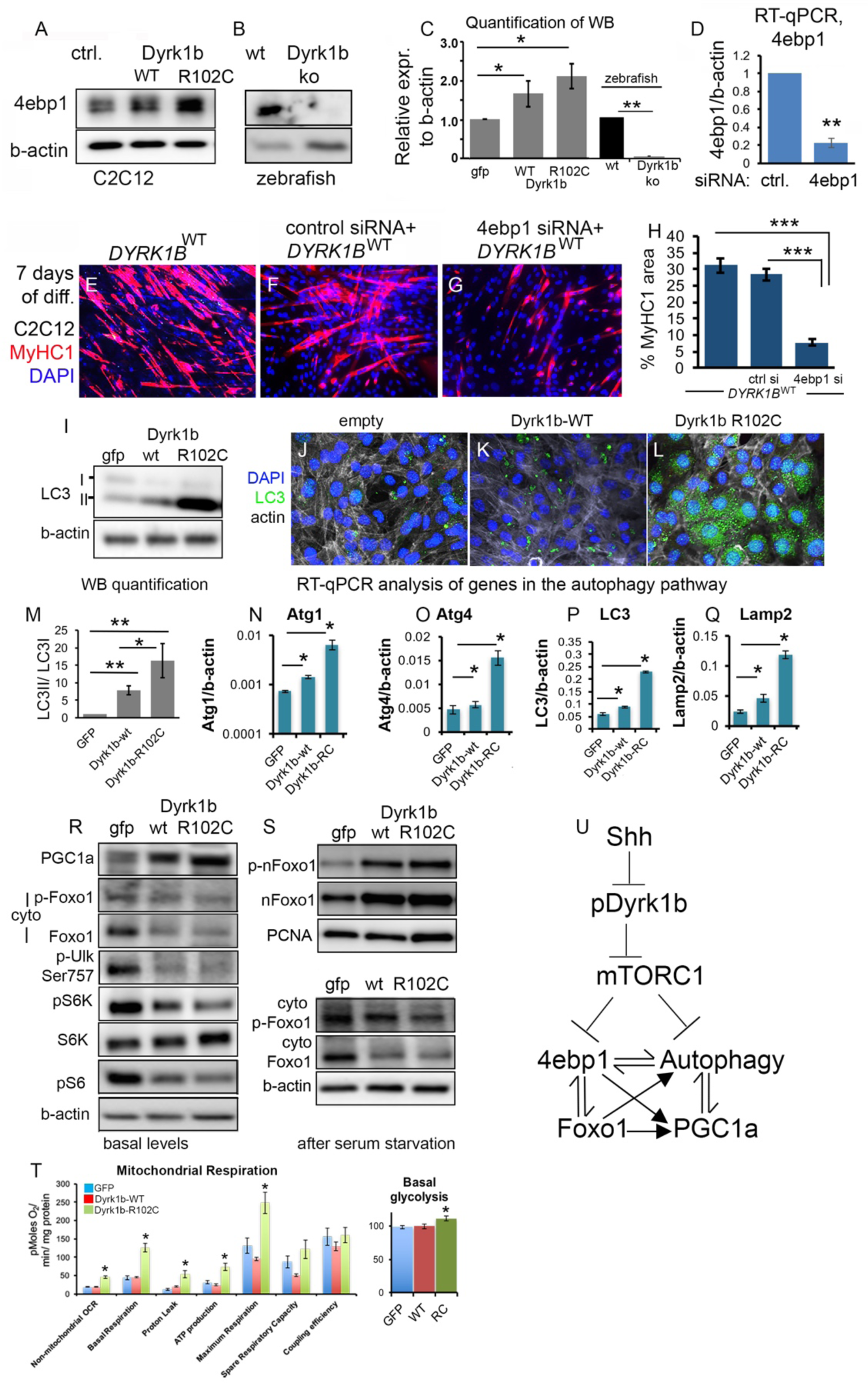
Dyrk1b inhibits mTORC1 signaling leading to increased 4ebp1, autophagy, PGC1a and nuclear Foxo1. **A-C** Western blot and quantification of (mean ± s.e.m) 4e-bp1 expression in C2C12 cells overexpressing empty vector, DYRK1B^WT^ or DYRK1B^R102C^. B WB showing 4e-bp1 expression in zebrafish wild type embryos or Dyrk1b knockout embryos. C Quantification of the blots shown in A-B. **D** RT-qPCR showing reduction of 4e-bp1 expression using 4ebp1siRNA compared to cells transfected with control siRNA. **E-G** IF against MyHC1 expression in C2C12 cells transfected with DYRK1B^WT^ alone, DYRK1B^WT^+ control siRNA, DYRK1B^WT^+ 4e-bp1 siRNA. The cells were transfected before the onset of differentiation and MyHC1 expression was examined on day7 of differentiation. **H** The percentage of MyHC1 area calculated for the conditions shown in panels E-G. **I-M** LC3 expression by WB (I, M) and IF (J-L) in the indicated backgrounds. **N-Q** RT-qPCR analysis of the proteins in the autophagic pathway in the C2C12 transfected with the indicated constructs. **R-S** WB displaying expression of indicated proteins collected from C2C12 cells that were transfected with the indicated constructs. The cells were transfected before the onset of differentiation and were collected on day 7 of differentiation in their basal state (R) or after serum-starvation (S). The top panels in S shows nuclear Foxo1 and the bottom panels shows cytoplasmic Foxo1. **T** Seahorse assay to examine mitochondrial respiration and glycolysis in C2C12 transfected with indicated constructs. **U** The model depicting Dyrk1b induced myofiber differentiation in C2C12 cells: during skeletal muscle differentiation in C2C12 cells, Shh inhibition promotes Dyrk1b phosphorylation and stimulates myoblast differentiation. Dyrk1b inhibits mTORC1 signaling that activates an interconnected network of increased translational inhibition by 4e-bp1, elevated autophagy, Foxo1 nuclear translocation and increased PGC1a.

Previous reports have suggested that 4e-bp1 activates a cross-regulatory interconnected network of autophagic pathways, PGC1a and Foxo1, during adult skeletal muscle homeostasis (Figure 5U) (62, 64-66). The autophagic flux was reported to be necessary and highly dynamic during C2C12 differentiation, peaking up at the beginning of C2C12 differentiation, and gradually tapering off towards the end of differentiation (Sin et al., 2016). This ensures timely degradation of ineffective mitochondria and production of mitochondria better suited for oxidative phosphorylation in myofibers. These events are in-turn co-incident with elevated mitochondrial biogenesis mediated by PGC1a (62, 64-66). Consistent with these reports, we found that elevated 4e-bp1 expression observed in DYRK1B^WT^ C2C12 cells is associated with increased autophagic flux as visualized by increased LC3II/LC3I ratio by WB (Fig.5 I, M, the experiment was done on n=4 biological replicates with identical results) and increased punctate LC3 immunostaining by the end of differentiation in C2C12 cells (Fig.5 J-L, n=2 biological replicates for each condition). The elevation was more pronounced in Dyrk1b^R102C^ relative to Dyrk1b^WT^ cells. Accordingly, the RNA expression of genes in the autophagic pathway such as Atg1, Atg4, LC3 and Lamp2b were increased in Dyrk1b^WT^ and even to a greater degree in Dyrk1b^R102C^ cells vs. GFP-transfected cells (Fig.5 N-Q). The increased autophagic flux was associated with elevated PGC1a expression in Dyrk1b^WT^ and Dyrk1b^R102C^ cells versus GFP-transfected cells (Fig.5R, n=2 biological replicates with identical results). This increased PGC1a expression in Dyrk1b^R102C^ myotubes correlated with significantly increased basal glycolysis and mitochondrial oxidation (Fig.5T, n=11 technical replicates for each transfection condition, n=2 independent experiments). Foxo1 has been previously associated with increasing autophagy and PGC1a expression (67, 68). We found that cytosolic Foxo1 levels were reduced (Fig.5R, S, n=2 independent experiments), while nuclear levels were elevated in Dyrk1b^WT^ and Dyrk1b^R102C^ cells versus GFP-transfected cells (Fig.5S). We next considered the possibility that Dyrk1b may suppress mTORC1 activity, which would encompass the molecular changes manifested as elevated 4ebp1, elevated autophagy, and elevated total Foxo1 levels (61). Accordingly, both pS6K and pS6 levels were reduced in Dyrk1b^WT^ and to a greater degree in Dyrk1b^R102C^ C2C12 versus GFP-transfected cells (Fig.5R, n=2 independent experiments). Consistent with these data, the proteomic analysis independently predicted activation of AMPK signaling, which counteracts mTOR pathways and induces autophagy (Fig.4C). Taken together, our findings reveal that Dyrk1b mediates suppression of mTORC1 signaling and subsequently activates an interconnected network of 4ebp1, autophagy, PGC1a, and Foxo1 (Fig.5U).

## DISCUSSION

By using a comprehensive *in vivo* and *in vitro* approaches, we demonstrate here the role of Dyrk1b as a key regulatory node that integrates multiple pathways involved in skeletal muscle development.

In vertebrates, skeletal muscle precursors in the segmental plate consists of two distinct adaxial and paraxial populations with unique fates. The adaxial cells are localized adjacent to the notochord and contribute to the slow muscle fibers while paraxial cells differentiate into fast-twitch fibers (Wolff et al., 2003, Reifers et al., 1998). Myogenesis in both cell populations is regulated by the master transcription factor (TF) MyoD. The *myoD* expression in adaxial cells is dependent on Shh signaling activity (69-71), while paraxial *myoD* positive cells are temporally and spatially regulated by Fgf8. The mechanisms that coordinate the activities of Fgf and Shh pathways was previously not known. Importantly, the downstream effectors of Fgf8 in myogenesis had not been explored. Using combined genetics and pharmacological approaches, we show here that Dyrk1b plays a critical role during early skeletal muscle specification by mediating FGF signaling in the paraxial domain and coordinating with Shh in the adaxial domain to induce *myoD* expression and initiate muscle specification. Accordingly, loss of Dyrk1b by gene editing in zebrafish led to loss of *myoD* in paraxial mesoderm while the overexpression of human Dyrk1b cDNA rescued loss of paraxial and adaxial *myoD* caused by Fgf and Shh inhibition, respectively. Similar interface of Shh and Fgf signaling is evident in the zebrafish retina where Fgf from dorsal forebrain specifies dorsal optic vesicle and Shh from ventral forebrain specifies ventral optic vesicle(72). Future work aims to determine how Dyrk1b may interact with these pathways during forebrain patterning, considering high expression of Dyrk1b in these domains and the cyclopic eye phenotype.

Further, our investigations of muscle differentiation in C2C12 mouse myoblasts revealed that Dyrk1b stimulates myogenic differentiation in a kinase-dependent manner. In this phase, Shh inhibition leads to excessive myogenic differentiation by promoting the Dyrk1b auto-phosphorylation. This finding shows again that Dyrk1b integrates Shh signaling, although in different ways, to advance the process of muscle differentiation into its completion. These functions of Dyrk1b make it an optimal candidate for development of therapeutics against sarcopenia.

Prior to our study very little was known about the downstream effectors of Dyrk1b. The proteomics analysis, supported by *in vivo* studies identified 4e-bp1 as one of the key targets of Dyrk1b. Strikingly, 4e-bp1 was found to be almost entirely absent in Dyrk1b-CRISPR knockout zebrafish embryos, indicating a cross-species conservation of 4e-bp1 regulation by Dyrk1b. Numerous reports indicate that 4e-bp1 activates a network of interconnected pathways to maintain skeletal muscle homeostasis by upregulation of PGC1a, Foxo1 and induction of autophagy in mice (62) and drosophila (65). Accordingly, C2C12 cells overexpressing Dyrk1b exhibited higher PGC1a protein levels, increased autophagy and elevated nuclear Foxo1 levels compared to the controls. Interestingly, the molecular changes orchestrated by elevated 4e-bp1 levels suggested reduced mTORC1 activity. Accordingly, mTORC1 activity was dramatically lower upon Dyrk1b overexpression in C2C12 cells. These results were also consistent with the increased activation of AMPK as an upstream regulator by proteomic analysis. Overall, these findings indicate the important role of DYRK1B in regulation of genes that play critical roles in skeletal muscle homeostasis.

One major strength of our study is its link to a human disease gene and hence the relevance of the discovery of Dyrk1b-dependent molecular networks to human skeletal muscle development and disease. The autophagy-PGC1a-4ebp1 network was considerably more activated in Dyrk1b^R102C^ vs, Dyrk1b^WT^ expressing cells, indicating a gain-of function effect as reported previously (10). The lack of muscle differentiation in Dyrk1b^R102C^ C2C12 cells indicates that a dynamic and fine-tuned regulation of the autophagy-PGC1a-4ebp1 network is required for proper myofiber differentiation. During the course of C2C12 differentiation, autophagic pathways are regulated in a dynamic fashion: LC3II/LC3I is elevated on day1 and reduces through the course of differentiation, concomitant with mitophagy and gradual generation of younger mitochondria with increased capacity for oxidative respiration of glucose (Sin et al., 2016). Functional characterization of Dyrk1b and its recognition as a key regulatory node for multiple myogenic pathways provides mechanistic insights into human skeletal muscle development and disease. Future genetic studies will determine whether mutations in gene encoding different components of Shh, Fgf and mTOR-4ebp1 pathways underlie disorders of musculoskeletal system.

In conclusion, our comprehensive characterization of Dyrk1b *in vivo* and *in vitro* identifies its role as a novel key regulatory node that integrates multiple pathways involved in myogenic differentiation and regeneration and a potential target for therapeutic interventions for the aging population with limited ability for muscle regeneration.

## MATERIALS AND METHODS

### Zebrafish lines and maintenance

Zebrafish strain WT^CF^ were raised and maintained according to protocols approved by the Yale University Institutional Animal Care and Use Committee (IACUC protocol number 2015-20070). Embryonic staging and husbandry were performed as described (73, 74).

### Whole-mount *in-situ* mRNA hybridization

Sense and antisense Dig-labeled probes were designed against zebrafish Dyrk1b transcripts. Antisense probe generated through PCR that amplified an exonic region of approximately 420 bp in the three Dyrk1b transcripts using the primers GCAGAGCCCACAAGCTCCGTCTCAGCAGC and<u>TAATACGACTCACTATAGGG</u>GGCCCAGGTGGAGGTTGGAGGAGCCGTAC. Second anti-sense probe spanning ∼ 550 bp was generated using <u>TAATACGACTCACTATAGGG</u>TCCCTGCGGCGGGTTGTTATTG and GGGACAGCAGCATGGGTCAGCT Sense probe was generated using GGCCCAGGTGGAGGTTGGAGGAGCCGTAC and <u>TAATACGACTCACTATAGGG</u>GCAGAGCCCACAAGCTCCGTCTCAGCAGC. (underlined region is the T7 promoter for In vitro transcription) These primers amplified exonic region of Dyrk1b and the product was cleaned using PCR purification kit (Qiagen)and cloned into pGEM®-T Easy Vector Systems (Promega). The sequence was confirmed through sanger sequencing. In vitro transcription was performed on the pcr product with Ampliscribe T7-flash kit with Dig RNA labeling mix from Roche(75). In situ hybridization with both the anti-sense probes yielded the same results.

### CRISPR/Cas9 gene editing, identification of On-target mutagenesis and Off-target analysis

gRNA sequence was generated through CHOPCHOP against the start codon and kinase domain. The gRNA sequence GCTGCTGGTCGGCCATGGAG<u>TGG</u> was designed to against the start codon and GAGAGCGGTAAAACCGGCTC<u>TGG</u> against the kinase domain respectively. PAM region is underlined and is excluded from the gRNA sequence. Generation of gRNA, Cas9 mRNA and zebrafish injections was done as described(36). To confirm the gRNA activity, genomic DNA was isolated using a DNeasy Blood and Tissue kit (Qiagen) from a clutch of 15-20 embryos at 24 hpf injected with gRNA and Cas9. 50 ng/µl or fin-clip of adult fish. This served as the template for pcr amplification of approximately 200-300bp region surrounding the intended target. Mutations were detected through a T7 endonuclease I assay (NEB)(76). To confirm the disruption of endogenous locus, the pcr product was cloned into pGEM®-T Easy Vector Systems (Promega) and sequence was analyzed after sanger sequencing. Once the mutation was detected, the rest of the embryos from the same clutch were raised to adulthood. PCR Fragment Analysis was also performed to analyze the indels generated through gRNA.

### Primer sequences for T7 endonuclease assay and pcr fragment analysis

Forward and reverse primer for amplifying region surrounding the start codon are TCCCTATGAAGGTGGAAGCA and CGGATGAGACAGTGAAACAAAGA respectively. Forward and reverse primer for amplifying region surrounding the kinase domain are TGTGTGTTTGTGCATGTCTTTC and CATCGTGTAACCGTACCTCATT respectively. The FAM were added to the forward primers for pcr fragment analysis.

### Off-target analysis

The gRNA sequence GCTGCTGGTCGGCCATGGAG was used as input in Chopchop (77-79)and CAS offfinder (CRISPR RGEN tools) (79) and no similar sequences were identified in the zebrafish genome that had 1, 2or 3 bp mismatches.

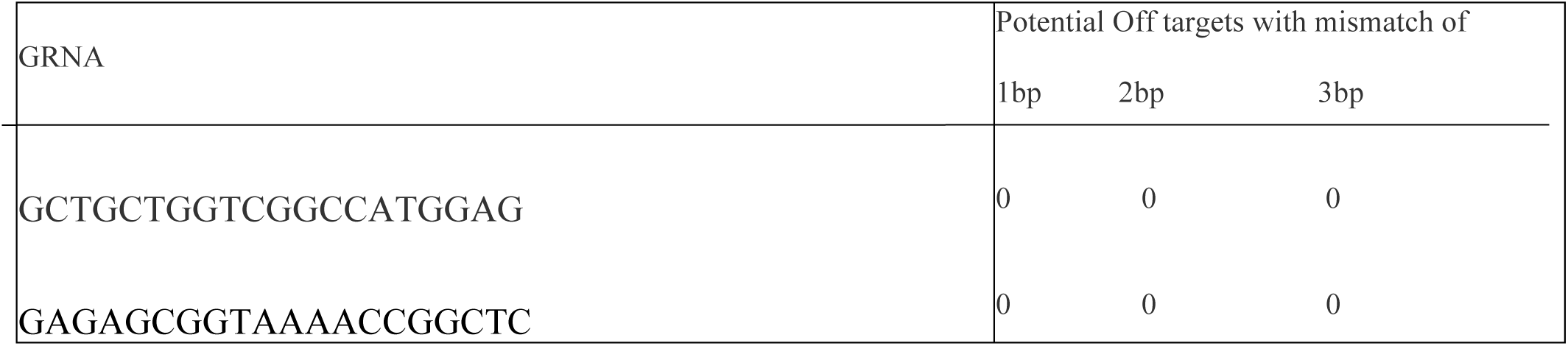

### Plasmids for *DYRK1B* overexpression, site directed mutagenesis for *DYRK1B*

The generation of plasmids for Dyrk1b and Dyrk1b ^R102C^ has been described (10). To generate Dyrk1b ^Y273F, K140R^, primers were designed with the desired point mutations flanked on either side by approx. 20 base pairs. The primers were PAGE purified and the template was PCR amplified by Turbo Taq DNA polymerase along with the mutated primers. The reactions with the forward and reverse primers were done separately, followed by Dpn1 digestion of parent plasmid, and transformation of plasmids into the Ultracompetent cells (Agilent). The plasmid was sequenced to verify the point mutation. Following primers were used for creating point mutations:

K140R: Forward: GACCCAGGAGCTTGTGGCCATCAGAATCATCAAGAACAAAAAGGC Reverse: GCCTTTTTGTTCTTGATGATTCTGATGGCCACAAGCTCCTGGGTC

Y273F: Forward: TGGCCAGAGGATCTACCAGTTCATCCAGAGCCGCTTCTACCG Reverse: CGGTAGAAGCGGCTCTGGATGAACTGGTAGATCCTCTGGCC

### mRNA synthesis and one-cell stage injections into zebrafish embryos of human Dyrk1b and Dyrk1b^R102C^ or Dyrk1b ^*Y273F, K140R*^

mRNA was synthesized after restricting digestion of the plasmid with XbaI and purified using phenol-chloroform extraction and ethanol precipitation. This purified product was in vitro transcribed with T7. The mRNA was purified using RNAeasy Mini kit (Qiagen, Catalog number 74104). Zebrafish embryos at the one-cell stage injected with 2 nanoliters of the solution containing 150 ng/μl of the mRNA and phenol red for visualization.

### Quantitative PCR of zebrafish embryos

Single cell stage embryos were injected with CRISPR/Cas9 against Dyrk1b locus or mRNA for Dyrk1b overexpression (o/e), Dyrk1b^R102C^ or Dyrk1b ^*Y273F, K140R*^. Embryos were raised to the required stage and RNA was isolated using 300μl of RLT-beta-mer (1 ml RLT buffer and 10 μl beta-mercaptoethanol). The embryos were dissociated by passing them through a syringe and needle (0.5mm X 22mm, catalog number 305124, BD). The embryos were then passed through a QIA shredder (Catalog number 79654) and purified using RNeasy Mini kit. cDNA synthesis was performed using iScript cDNA synthesis kit (Bio-Rad, 1708891). qPCR was performed using iTaq Universal SYBR Green Supermix (Bio-Rad 172-5121). All reactions were done in triplicates and quantified in relation to beta-actin.

Primers for qPCR in zebrafish:

<u>Genes</u>

*myoD* Forward: ACTATCCCGTTCTGGAACATTAC

Reverse: GTCATAGCTGTTCCGTCTTCTC

*dyrk1b* Forward: AGCAGGTGCTGTCAGATA

Reverse: GATGAGATCGACTGAGAGTTT

*pea3* Forward: CCCGCCTGTGAAGATTAAGAA

Reverse: CTCATAGGCACTGGCGTAAA

*sprouty* Forward: CGTGCATGTGTTTGGTGAAG

Reverse: ACTGCGGATGCGATAATGAA

*sef* Forward: CACCTTCGGCTTCTCCATTTA

Reverse: CATACGTTCCTGGAGTGACATC

*b actin* Forward: CGAGCTGTCTTCCCATCA

Reverse: TCACCAACGTAGCTGTCTTTCTG

### Differentiation and transfection of C2C12 cells

The C2C12 cells were maintained in 10%FBS+ DMEM+1X Pen-Strep. The differentiation was induced by shifting the media to 2%horse serum+ DMEM+ 1X Pen-Strep. The cell culture media was changed every day until the end of differentiation at day7. The plasmids containing human wildtype *DYRK1B, DYRK1B*^*R102C*^, *DYRK1B*^*Y273F, K140R*^ or GFP were transfected before ensuing the differentiation, using lipofectamine 3000 (Invitrogen) using manufacturers’ guidelines with a slight modification: The DNA-lipofectamine mix was added to C2C12 cells in suspension post trypsin treatment. For immunostaining, the transfected cells were cultured on glass coverslips and subjected to the standard protocols. For western blot, the cells were harvested in cell lysis buffer+protease inhibitors (Cell Signaling). The cell fractionation into nuclear and cytoplasmic fractions was done using NE-PER extraction method (Thermo Fisher Scientific).

### Inhibition of Shh and Fgf pathways

Inhibition of Shh was done through Cyclopamine (LC laboratories) as described (80).

Inhibition of FGF8 was done through incubation of embryos in the drug SU5402, dilution of the drug was described(81). The cyclopamine and SU5402 treatments were added at a final concentration of 1µM from a stock of 10mM for each. Equivalent amount of DMSO was added to control cultures. The treatment was done for 7 days in the differentiation media and the media was changed every day.

### Western Blot for zebrafish embryos

The zebrafish lysates at 14hpf and 24hpf were prepared in 0.063M Tris, pH 6.8, 10% glycerol, 0.5ml Beta-mercaptoethanol (in a total of 10mls), 3.5% SDS (zebrafish book). For each sample, 40-50 embryos were pooled together. The lysates were heated at 95°C for 10 minutes prior to running the SDS-PAGE gel. Western blot was carried by standard procedures. All quantifications were done normalized to beta-actin.

### Antibodies for Western Blot and Immunostaining

The F59 and A4.1025 (MyHC1) antibodies were obtained from Developmental Studies Hybridoma Bank. Gli1 (Abcam, ab49314), Gli2a (Genetex, GTX128280), Gli3 (Genetex, GTX26050), MyoD (Santa Cruz, sc-302, Zfin), pHIPK2 (PA513045, Life technologies), beta actin (Abcam, ab8227), LC3 (Cell signaling, 2775)Pgc1a (Novus Biological NBP1-04676) antibodies were all selected and purchased based on their prior verifications. The western blots were quantified by ImageJ.

### Proteomics: Tissue processing for proteomics

The tissues were provided to the Yale proteomics core and suspended in 500uL RIPA buffer contains protease (Pierce; 100X:cat#87786) and phosphatase (Pierce; 100X:cat#78420) inhibitor cocktail. Three bursts (10% amp) followed by two bursts (15% amp) of sonication for 15 seconds each were carried out to burst the cells open. Suspension was centrifuged at 14K rpm for 10 minutes, and 150uL of the supernatant was removed. Chloroform:methanol:water protein precipitation was performed, and dried protein pellet was resuspended in 100uL 8M urea containing 400mM ABC, reduced with DTT alkylated with iodoacetamide, and dual enzymatic digestion with LysC and trypsin (carried out at 37 C for overnight and 6 hrs., respectively). Digestion was quenched (with 0.1% formic acid) during the de-salting step with C_18_ MacroSpin columns (The Nest Group). The effluents from the de-salting step were dried and re-dissolved in 5 µl 70% FA and 35 µl 0.1% TFA. An aliquot was removed from each sample to determine peptide concentrations via Nanodrop, and diluted to 0.05 µg/µl with 0.1% TFA. 1:10 dilution of 10X Pierce Retention Time Calibration Mixture (Pierce; Cat#88321) was added to each sample prior to injecting on the UPLC Q-Exactive Plus mass spectrometer to check for retention time variability the normalization during LFQ data analysis.

<u>Label-Free Quantitation (LFQ)</u> LC MS/MS data collection was performed on a Thermo Scientific Q-Exactive Plus Mass spectrometer connected to a Waters nanoACQUITY UPLC system equipped with a Waters Symmetry® C18 180 µm × 20 mm trap column and a 1.7-µm, 75 µm × 250 mm nanoACQUITY UPLC column (35°C). 5 µl of each digests (in triplicates) at 0.05 µg/µl concentration was injected in block randomized order. To ensure a high level of identification and quantitation integrity, a resolution of 60,000 was utilized for MS and 15 MS/MS spectra was acquired per MS scan using HCD. All MS (Profile) and MS/MS (centroid) peaks were detected in the Orbitrap. Trapping was carried out for 3 min at 5 µl/min in 99% Buffer A (0.1% FA in water) and 1% Buffer B [(0.075% FA in acetonitrile (ACN)] prior to eluting with linear gradients that will reach 30% B at 140 min, 40% B at 155 min, and 85% B at 160 min. Two blanks (1st 100% ACN, 2nd Buffer A) will follow each injection to ensure against sample carry over. The LC-MS/MS data was processed with Progenesis QI software (Nonlinear Dynamics, version 2.4) with protein identification carried out using in-house Mascot search engine (2.4). The Progenesis QI software performs chromatographic/spectral alignment (one run is chosen as a reference for alignment of all other data files to), mass spectral peak picking and filtering (ion signal must satisfy the 3 times standard deviation of the noise), and quantitation of peptides and proteins. A normalization factor for each run was calculated to account for differences in sample load between injections as well as differences in ionization. The normalization factor was determined by calculating a quantitative abundance ratio between the reference run and the run being normalized, with the assumption being that most proteins/peptides are not changing in the experiment so the quantitative value should equal 1. The experimental design was setup to group multiple injections (technical and biological replicates) from each run into each comparison sets. The algorithm then calculates the tabulated raw and normalized abundances, Anova *p* values for each feature in the data set. The MS/MS spectra was exported as .mgf (Mascot generic files) for database searching. Mascot Distiller was used to generate peak lists, and the Mascot search algorithm was used for searching against the Swiss Protein database with taxonomy restricted to *Mus musculus*; and carbamidomethyl (Cys), oxidation (Met), Phospho (Ser, Thr, Tyr), and Deamidation (Asn, Gln), Acetyl (Lys) were entered as variable modifications. Two missed tryptic cleavages were allowed, precursor mass tolerance was set to 10ppm, and fragment mass tolerance was set to 0.02 Da. The significance threshold was set based on a False Discovery Rate (FDR) of 1%. The Mascot search results was exported as .xml files and then imported into the processed dataset in Progenesis QI software where peptides ID’ed were synched with the corresponding quantified features and their corresponding abundances. Protein abundances (requiring at least 2 unique peptides) were then calculated from the sum of all unique normalized peptide ions for a specific protein on each run.

A label free quantitation approach utilizing Progenesis QI (Nonlinear Dynamics) similar to that found in (82) was utilized to obtain quantitative information on peptides and proteins which can be referenced in the method.

Three independent samples from each genotype: empty vector and DYRK1B-wt transfected C2C12 cells were analyzed for proteomics analysis. The cells were serum starved prior to submission for proteomics analysis. The fold changes corresponding to each protein were calculated and subjected to the following filters: unique peptides >2, p-value ≤ 0.05, 2-tailed unpaired ttest, FDR ≤ 0.05, Benjamini Hochenberg (B-H) correction. The significantly changed proteins were used as inputs for Ingenuity Pathway Analysis (IPA, Version 03-13, Qiagen). The IPA predicted the altered pathways, molecular and cellular functions, and potential upstream regulators which are reported. For the latter, the data was filtered using p-value ≤ 0.05 and -2<z-score <2. The hierarchical clustering of the variables and the heat-maps for differentially expressed proteins were created by Qlucore Omics Explorer 3.4.

### Statistical information

All statistical analyses were carried out using 2 tailed Student’s ttest for pairwise and Anova one-way analysis for multiple comparisons using GraphPad Prism 6 Project software (GraphPad). P ≤ 0.05 was used as statistical threshold.

Dyrk1b: Dual-specificity Tyrosine-regulated kinase 1B
SO: sarcopenic obesity
MetS: metabolic syndrome
myoD1: myoblast determination protein 1
Fgf: Fibroblast growth factor
Shh: Sonic Hedgehog
mTORC1: mechanistic target of rapamycin C1
4e-bp1: Eukaryotic translation initiation factor 4E (eIF4E)-binding protein 1

## ACKNOWLEDGEMENTS

The authors thank Dr. Antonio J. Giraldez, for the zebrafish in situ hybridization probe and Dr. Arne C. Lekven and Dr. Bruce B. Riley for gifts of cDNAs for zebrafish in situ probes and plasmids. We would also like to thank Ms. Meredith S. Cavanaugh, Aquatics Facility Manager for the fish care. We also thank Developmental Hybridoma Bank for the antibodies. This work was supported by the NIH grant RHL135767A to A.M.

## AUTHOR CONTRIBUTIONS

N.B. (**Neha Bhat Ph. D**) and A.N., (**Anand Narayanan Ph. D**) were primarily responsible for performing the experiments and the analysis and helped with the study, design and writing of the manuscript

A.M. (**Arya Mani M.D.**) designed the study and oversaw its implementation, supervised all aspects of the project from performing the experiments to the analysis of all data and wrote the manuscript.

## DECLARATION OF INTERESTS

The authors declare no competing interests

---

**Table S1.** Homozygous and heterozygous embryos from a clutch of Dyrk1b^+/-^ intercross

| Number of embryos in a clutch of ~ 100 F2 embryos |  |  |  |
| --- | --- | --- | --- |
| Homozygous | Heterozygous | Wild type | Total number of embryos |
| 23 | 48 | 24 | 95 |
| 26 | 40 | 32 | 98 |
| 30 | 35 | 40 | 105 |

